# Genome-wide CRISPR knockout screen reveals membrane tethering complexes EARP and GARP important for Bovine Herpes Virus Type 1 replication

**DOI:** 10.1101/2020.06.17.155788

**Authors:** Wenfang S. Tan, Enguang Rong, Inga Dry, Simon G. Lillico, Andy Law, Christopher B.A. Whitelaw, Robert G. Dalziel

## Abstract

We produced a genome wide CRISPR knockout library, btCRISPRko.v1, targeting all protein coding genes in the cattle genome and used it to identify host genes important for Bovine Herpes Virus Type 1 (BHV-1) replication. By infecting library transduced MDBK cells with a GFP tagged BHV-1 virus and FACS sorting them based on their GFP intensity, we identified a list of pro-viral and anti-viral candidate host genes that might affect various aspects of the virus biology, such as cell entry, RNA transcription and viral protein trafficking. Among them were VPS51, VPS52 and VPS53 that encode for subunits of two membrane tethering complexes EARP and GARP. Simultaneous loss of both complexes in MDBKs resulted in a significant reduction in the production of infectious cell free BHV-1 virions, suggesting the vital roles they play during capsid re-envelopment with endocytosed membrane tubules prior to endosomal recycling mediated cellular egress. We also observed potential capsid retention and aggregation in the nuclei of these cells, indicating that they might also indirectly affect capsid egress from the nucleus. The btCRISPRko.v1 library generated here greatly expanded our capability in BHV-1 related host gene discovery; we hope it will facilitate efforts intended to study interactions between the host and other pathogens in cattle and also basic host cell biology.

## Introduction

Bovine Herpes Virus Type 1(BHV-1) causes major economic loss through multiple disease manifestations such as infectious bovine rhinotracheitis (IBR), infectious pustular vulvovaginitis, balanoposthitis and abortions in cows^1^. The virus is endemic in Ireland and the UK, with 80% of herds seropositive for infections^2–4^. Following successful clearance of acute infection, BHV-1 establishes lifelong latency in sensory neurons via neuronal retrograde transport. Stressful life events such as a pregnancy, co-mingling and transporting during extreme weather can trigger reactivation to induce clinical diseases. Vaccination can be used to control IBR; however, a recent study showed that strains isolated from respiratory cases and aborted fetuses corresponded to modified live vaccine strains^5^. Improved knowledge in host-pathogen interactions is needed to develop better disease control and resistance strategies.

Like other alpha-herpesvirus, BHV-1 replicates its dsDNA genome in the host nucleus but matures into infectious particles in the cytoplasm. To initiate a lytic cycle of infection, envelope glycoprotein gC of the virus interacts with cell surface Heparan Sulfate proteoglycans^6,7^, enabling close contact and binding between other glycoproteins and cell surface proteins. The binding of gD to the poliovirus receptor (PVR) and nectin-1^8–10^ and interaction between gB or gH/gL with putative receptor paired immunoglobulin-like type 2 receptor α or PILRα^11–14^ brings the virus closer to the cellular membrane and catalyzes cell penetrance by membrane fusion or endocytosis^15^. Upon entry, the viral capsid and tegument proteins are released into the cytoplasm to carry on infection. While the tegument proteins execute intricate plans of usurping or mitigating specific host factors for tight control of the environment in- and outside the nucleus, the capsid and associate viral tegument proteins are transported to the nucleus along the microtubule system, docks at the nuclear pore and injects the carrier genome into the nucleus for DNA replication and three stages of viral transcription, immediate early (IE), early (E) and Late (L)^16^. The transcripts are shuttled into the cytoplasm for translation and capsid subunits are transported back to the nucleus for capsid assembly and genome packaging prior to nuclear egress.

According to the envelopment-deenvelopment-reenvelopment model, the capsid of alpha-herpesviruses emerges from the nucleus mainly through primary envelopment at the inner nuclear membrane (NM) and deenvelopment at the outer NM^17–19^. Studies in Human Simplex Virus Type 1(HSV-1) found that before re-envelopment, virus surface glycoproteins are first secreted onto the plasma membrane (PM)^20,21^ and retrieved back to the cytoplasm via Rab5 GTPase dependent endocytosis^21,22^. The capsid regains membrane decorated with these glycoproteins from a cytoplasmic source prior to exocytosis; the event of secondary envelopment is a complex process and remains a contended topic in alpha-herpesvirus research. Previous studies on HSV-1 suggest that prior to egress via endosomal recycling^20^, membrane tubules of both early endosome^20^ and the trans-Golgi Network (TGN)^18,23,24^ origin can be used to wrap newly synthesized capsids into infectious virions, although more recently studies have cast strong doubt on TGN being a major alternative membrane source for re-envelopment^20,21^. Comparative ultra-structural studies between the HSV-1 and BHV-1 demonstrate that BHV-1 assembly utilizes recently endocytosed material for envelopment in a manner analogous to that of HSV-1, while side-by-side proteomic analysis further confirmed similar compositions for HSV-1^25^. It is discovered that the post TGN re-envelopment of HSV-1 capsids is Rab5 and Rab11 dependent^20,22^, but the detailed mechanisms linking the glycoprotein containing vesicles such as early endosomes (EE) to the recycling endosomes during secondary envelopment are still poorly understood.

Recent developments in genome editing technologies, such as TALENs and CRISPR/Cas9, offer unprecedented opportunities to study host-pathogen interactions such as virus envelopment and egress. CRISPR/Cas9 is an endonuclease and RNA complex that binds to target DNA via base pairing between the guide and target DNA contiguous with a protospacer adjacent motif (PAM) and executes cutting to induce gene knockout^26–28^. It is straightforward to reprogram CRISPR/Cas9 to target almost any genomic sequences, simply by altering the 20bp of guide sequence to be complementary to intended DNA targets. This unique feature enables high throughput targeting, disrupting many genes in parallel by delivering large number of guides in a pooled or arrayed manner. This is exemplified by genome wide CRISPR knockout screens that identified novel genes involved in infections from viruses such as flaviviruses, human influenza viruses and HIV^29–32^. During implementation, a large number of CRISPRs are deployed and the CRISPR serves two purposes once integrated in a cell, to inactivate the target gene and barcode the targeted cell for tracking. As the CRISPR is delivered via a lentivirus vector the sequence is integrated into the host chromosome; the range of CRISPRs present in a cell population determined by sequencing allow the identification of genes that are preferentially depleted or enriched in cells with a phenotype of interest.

To harness the power of genome wide CRISPR screens for gene discovery in cattle, we aimed to create a knockout library that would target all annotated coding genes in the cow genome. Here we present the process and success of generating this library, named btCRISPRko.v1, and our experience in using it to gain new insights into cellular factors important for active BHV-1 infection. We identified more than 150 such cellular factors, demonstrating the power of CRISPR screens in farm animal pathogen research. We concentrated our efforts on some of the top and novel hits to validate and investigate further, VPS50 to VPS54, five genes that encode subunits of two membrane tethering complexes EARP and GARP. Below we will detail our experiments and results that suggest crucial roles of these complexes play in secondary envelopment of BHV-1 and other alpha-herpesviruses. The knockout library btCRISPRko.v1 is readily available to study other pathogens in cattle and we hope it will become useful resource to the wider infectious disease research community.

## Results

### Overview of the genome wide CRISPR knockout screens

To identify host genes with candidate anti- or pro-viral roles, we need to isolate cells with differential responses to BHV-1 infection due to CRISPR induced loss of gene function, and a robust assay to harvest cells with relevant and distinct phenotypes from the screen is crucial. From our colleagues at ETH^33^, we obtained a recombinant BHV-1 virus strain with GFP fused to the C terminus of VP26, a product of the UL35 gene. This strain is based on BHV-1.1(strain Jura), the most prevalent respiratory form around the world^34^. A full cycle of BHV-1 infection takes around eight hours to complete (unpublished data); VP26 is a minor capsid protein synthesized during the late stage of infection, usually between four to six hours post infection in our hands. After infection of library transduced cells (**Fig. 1**), we can collect fractions of live cells using FACS sort based on cellular GFP intensity, a signal that reflects robustness of virus replication. Genomic DNA samples from these fractions can then be used for PCR to amplify the CRISPR regions for NGS and the copy numbers of all CRISPRs are obtained by counting matching reads. Due to the disruptive nature of guides, enrichment of guides leads to depletion of target genes and vice visa. With purposely designed data analysis packages such as MAGeCK^35^, we can compare CRISPR copy numbers between fractions, to identify genes with significantly enriched or depleted guides in these fractions relevant to the controls, and assign anti- or pro-viral roles to them prior to validation.

**Figure 1.**
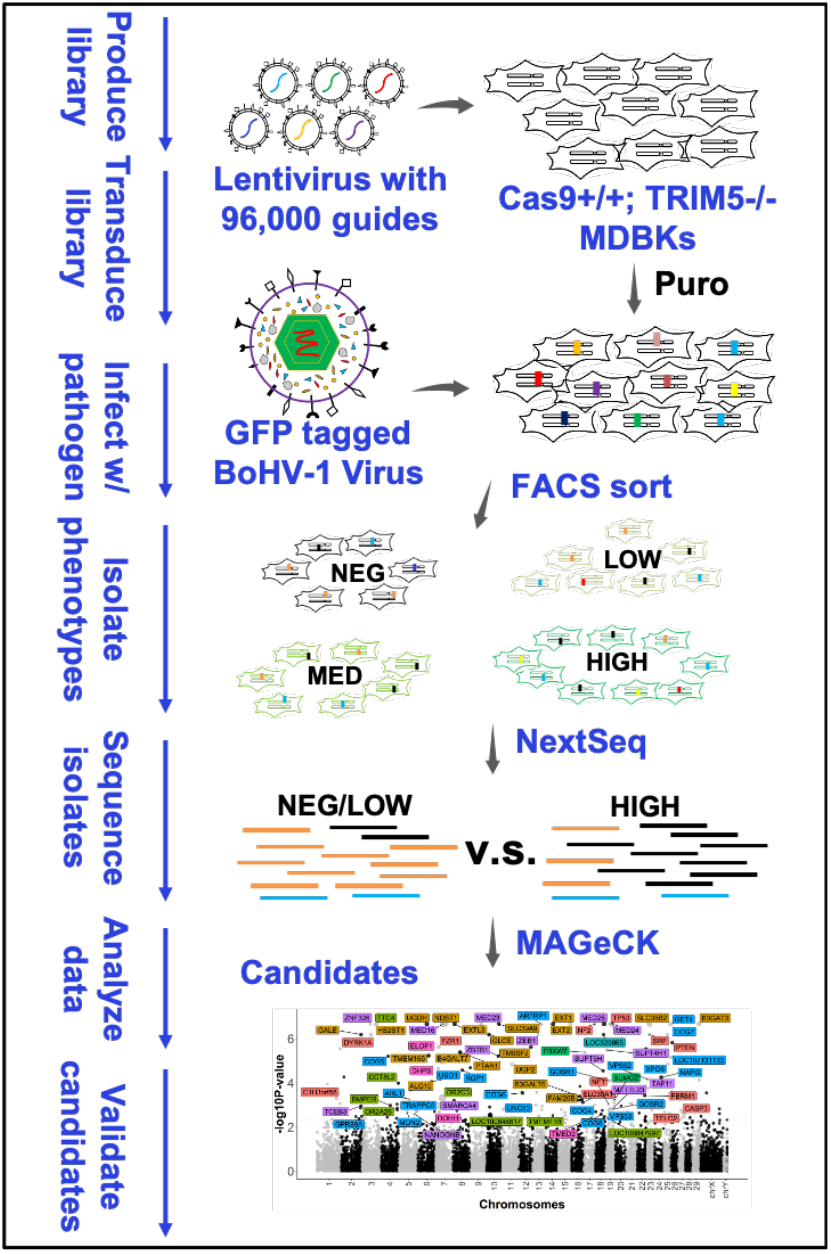
Overview of a CRISPR knockout screen for host gene discovery using BHV-1 as a model. A genome wide CRISPR knockout library is packaged into lentivirus and transduced into a highly transducible cow cell line MDBK engineered to express Cas9 from the rosa26 locus. After Puromycin selection, the transduced cells are challenged by a GFP tagged BHV-1 virus and FACS sorted based on their GFP intensity, an indicator of virus replication. This procedure separates the live cells into four sub-populations, GFP Negative, Low, Medium, and High. Genome DNA is isolated from these sub-populations and used as template for PCR to amplify the inserted CRISPR regions. The PCR products are sequenced by Illumina NextSeq and trimmed reads matching each guide in the library are counted. With MAGeCK, read counts from each sub-population are compared to identify enriched or depleted guides and their target genes. These genes are asigned pro-viral or anti-viral roles dependent on context, and are validated and further investigated.

### A highly transducible MDBK cell line with stable expression of Cas9

A highly susceptible cell line with uniform levels of Cas9 expression will greatly facilitate the screen and increase statistical power. By TALEN mRNA (**Fig. 2A**) mediated targeting of an EF1a promoter driven Cas9 expression cassette to the bovine rosa26 locus, we generated homozygous MDBK clones (**Fig. 2B**) with biallelic expression of Cas9. Unfortunately, lentiviral transduction efficiency in these MDBK clones was too low to deliver a genome wide CRISPR library cost-effectively for satisfactory coverage (**Fig. S1**). TRIM5a is an Interferon stimulated gene that has been shown to inhibit multiple RNA viruses including HIV-1 based lentivirus in cow cell lines^36–38^. Thus, we derived new clones with bi-allelic TRIM5a knockout and homozygous Cas9 expression after transfecting the mixture of cells recovered from the previous transfection with TALEN mRNA targeting TRIM5a (**Fig. 2C, Table S5**). We named them TRIM5a −/−; Cas9+/+, and the transduction efficiency in these clones increased eight fold (**Fig. 2D**).We compared BHV-1 infection in wild type (wt), Cas9+/+, and TRIM5a −/−; Cas9+/+ MDBK clones and observed no significant difference in plaque numbers and sizes (**Fig. S2,S3,S4**). These results indicate that TRIM5a does not interfere with BHV-1 replication and the modified cell lines support BHV-1 infection equally as the wt. We thus conducted all our BHV-1 screens using these TRIM5a −/−; Cas9+/+ MDBK cells and used Cas9+/+ clone #3 (**Fig. 2B**) for subsequent candidate validations.

**Figure 2.**
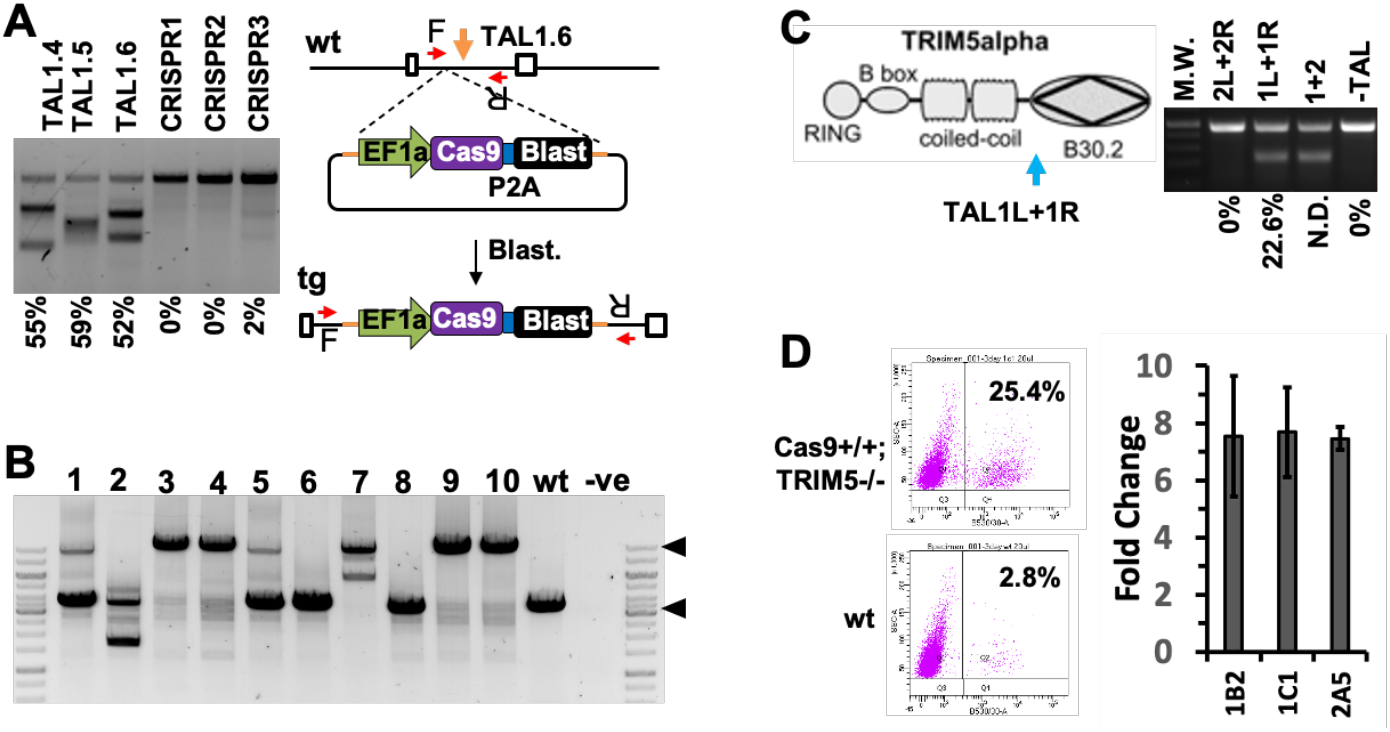
Production of a highly transducible Cas9 expressing MDBK cell line for the knockout screen. **A.** Knock-in (KI) Cas9 expression cassette to the rosa26 locus in MDBK cells. TALEN mRNA pair TAL1.6 cutting in first intron of the rosa26 locus (orange arrow) was co-transfected with plasmid containing the EF1a promoter driven Cas9-2a-Blasticidin expression cassette flanked by homology arms (HA, orange line in right panel). Single cell clones were isolated by dilution cloning after Blasticidin selection and genotyped using PCR primer F+R (red arrows). Percentages shown are estimated cutting efficiency based on ImageJ densitometry analysis. **B.** Genotyping results from ten clones. PCR amplicons from the wt (4Kb) and tg (targeted, 11Kb) alleles are indicated by the black triangles. Clones 1,5 are heterozygous targeted (Cas9+/wt), 3,4,9,10 are homozygotes (Cas9+/+) and 2,6,8 are not targeted. **C.** TRIM5a KO to enhance lentiviral transduction efficiency in MDBKs. TALENS designed to cut upstream of the B30.2 domain (blue arrow) to induce protein truncation. The population of cells after Blasticidin selection from A. was transfected with TAL1L+1R mRNA and clones were isolated and genotyped for both Cas9 KI and TRIM5a KO by PCR and sequencing. Cutting efficiency shown was based on TIDE analysis. N.D.: not determined. **D.** Three clones homozygous for Cas9 KI and TRIM5a KO (Cas9+/+; TRIM5−/−) were transduced with GFP lentivirus and the transduction efficiency was measured by FACS (n=3), bar chart represents fold changes relative to wt cells.

### CRISPR library designed with optimal targeting efficiency and specificity

To make the library broadly applicable, we combined the UMD3.1.1 assembly with the Y chromosome from btau5.0.1 and extracted all RefSeq coding transcripts. For each gene, sequences shared among isoforms were scanned for 20bp sequences followed by the Cas9 PAM NGG (**Fig. 3A**), and only CRISPRs that met our initial design criteria were kept (**Supplementary Materials and Methods**). Each CRISPR was estimated for the on-target efficiency using Azimuth 2.0^39^ and then aligned back to the genome using BWA^40^, to identify 20bp off-targets with up to three mismatches. 4-5 guides were picked by the following rules: 1) they have the highest estimated on-target efficiency; 2) they cut close to the 5’ of the CDS; and 3) they have the least off-targets, especially those that reside in coding sequences. The final library (**Data file 1**) contains 96,000 guides targeting 21,165 genes in total (**Table S1**) and 93.3% of the guides cut in the first half of common sequences among isoforms (**Fig. 3D**). The medium cutting efficiency is 0.67 with 0.61 and 0.72 at the 25% and 75% quantiles (**Fig. 3B**). 73% of all guides have no predicted off-targets residing in coding sequences with a CFD score >0.2(**Fig. 3C**)^39^. The off-targets residing in other regions of the genome were also controlled, as 73% of all guides have less than three off-targets with CFD >0.2 outside the coding sequences (**Fig. 3C**). Prior to library synthesis, we validated the design by testing guides against a few host genes being involved in alphaherpesvirus replication as indicated by literature^41–44^. All guides tested induced indels at the target sites after four days of Puromycin selection, with cutting efficiencies up to 79% (**Fig. S5**).

**Figure 3.**
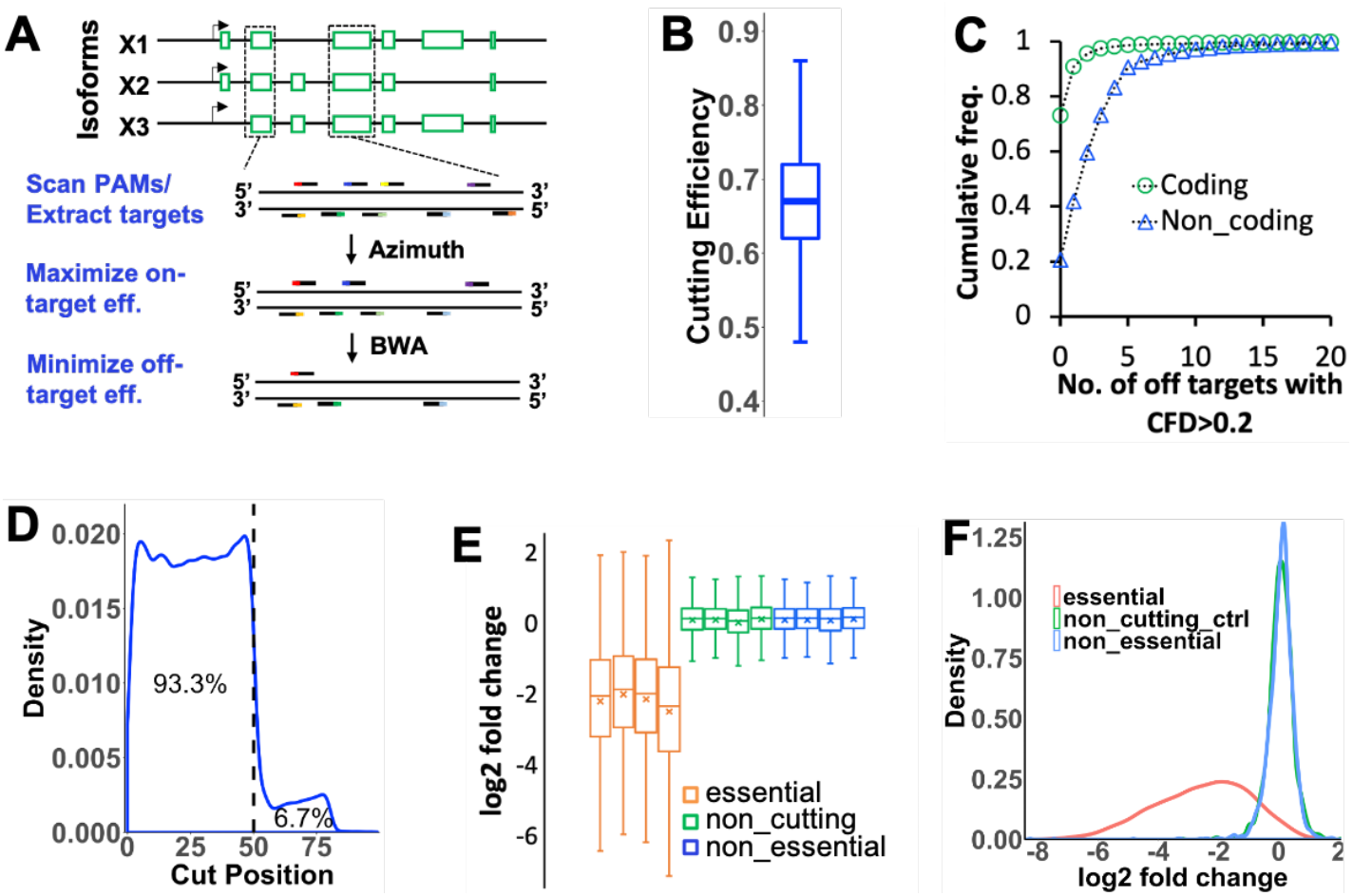
Lib design, statistics and performance assurance. **A.** pipeline developed for Genome-wide CRISPRko library design; **B.** Distribution of cutting efficiency estimated by Azimuth 2.0 for all guides in the library (**Data file 1**); **C.** Cumulative frequency of guides that have zero to 20 off-targets with CFD scores >= 0.2 within coding or non-coding sequences (**Data file 1**); **D.** Distribution of cutting sites of all 94,000 guides relative to the 5’ of common coding sequences across the genome (**Data file 1**); **E.** Distribution of log2 fold changes (l2fc) of guides targeting CEG2.0 genes(orange), non-essential genes(blue) and non-targeting gRNA controls(green) in cells transduced with the lentiviral K2g5 library relative to the copy numbers in the plasmid library (**Data file 2**). Results are based on four biological repeats from three independent transductions, cells from one transduction was used twice. **F.** Distribution of average l2fc for the three groups of guides from the four repeats(**Data file 2**).

### Library production and performance assurance in MDBK cells

Two sgRNA scaffolds are widely used, the original scaffold from the Broad Institute^26^ (referred as g2 here) and the optimized version by Chen et al^45^ (referred as g5). To determine which one delivers superior performance in MDBKs, we cloned the same oligo pool into two otherwise identical lentivirus vectors with the different scaffolds^46,47^ (**Fig. S6**; **Table S2**). We also generated two PiggyBac^48^ libraries bearing the g2 or g5 scaffold for screens to be conducted in hard to transduce cells (**Fig. S7; Table S2**). We transduced the lentiviral libraries, K2g2 and K2g5, into TRIM5a −/−; Cas9+/+ MDBKs, and after Puromycin selection sequenced them to study gRNA distributions (**Fig. S8, S9; Supplementary Materials and Methods**). We saw a higher depletion rate of guides targeting core essential genes(CEG2.0)^49^ in K2g5 containing cells than those in K2g2 cells (**Fig. S10, Data file S1**). The CEG2.0 is a curated list of fitness genes essential for cultured mammalian cells; the dropout rate of CEG2.0 targeting guides is a good indicator of library performance because CEG2.0 targeting CRISPRs with better activity are eliminated faster from the cellular pool. Thus, we chose the K2g5 library transduced cells for our BHV-1 screens as the bigger shift observed points to superior library performance. Re-sequencing of the same K2g5 cells with higher sequencing depth confirmed the shift compared to those targeting non-essential genes (**Fig. 3E, F, Data file 2**), and also the relative unaltered representation of non-cutting controls, indicating good off-targeting control measures implemented during the library design. We also detected the presence of 99.9% of guides in the K2g5 plasmid library and 96.01% of the reads mapped to the library without any mismatch. The library has a GiniIndex of 0.1258 assessed by MAGeCK (**Data file 2**)^35^, indicating good gRNA distribution. Based on these results, the quality and performance of the K2g5 library is extremely satisfactory.

### CRISPR knockout screen identifies pro-viral candidates involved in many aspects of virus biology

We conducted two rounds of screens by challenging K2g5 transduced cells with the tagged virus^33^ (**Fig. 1, Fig. S11**). During the 1^st^ screen, we infected cells at a MOI =2 and 10 hours post infection (10 h.p.i) sorted live cells into four fractions, GFP Negative, GFP Low, GFP Medium, and GFP High (**Fig. S12**, **Table S3**). The experiment was repeated three times and after sequencing and data analyses, we identified many genes with well-established roles in BHV-1 replication and others with undocumented roles (**Data files 2,4**). Encouraged by these results, we performed a 2^nd^ screen by infecting cells at the same MOI but FACS sorting them at 8 hpi instead. The earlier time point allowed more stringent gatings (**Fig. S12**) during sorting to isolate more extreme phenotypes and enable more cells to be recovered from FACS (**Table S4**). Reassuringly, this screen recapitulated the majority of candidates highlighted in the 1^st^ screen, along with many other new candidates (**Fig. 4A**, **Data files 3,4**).

**Figure 4.**
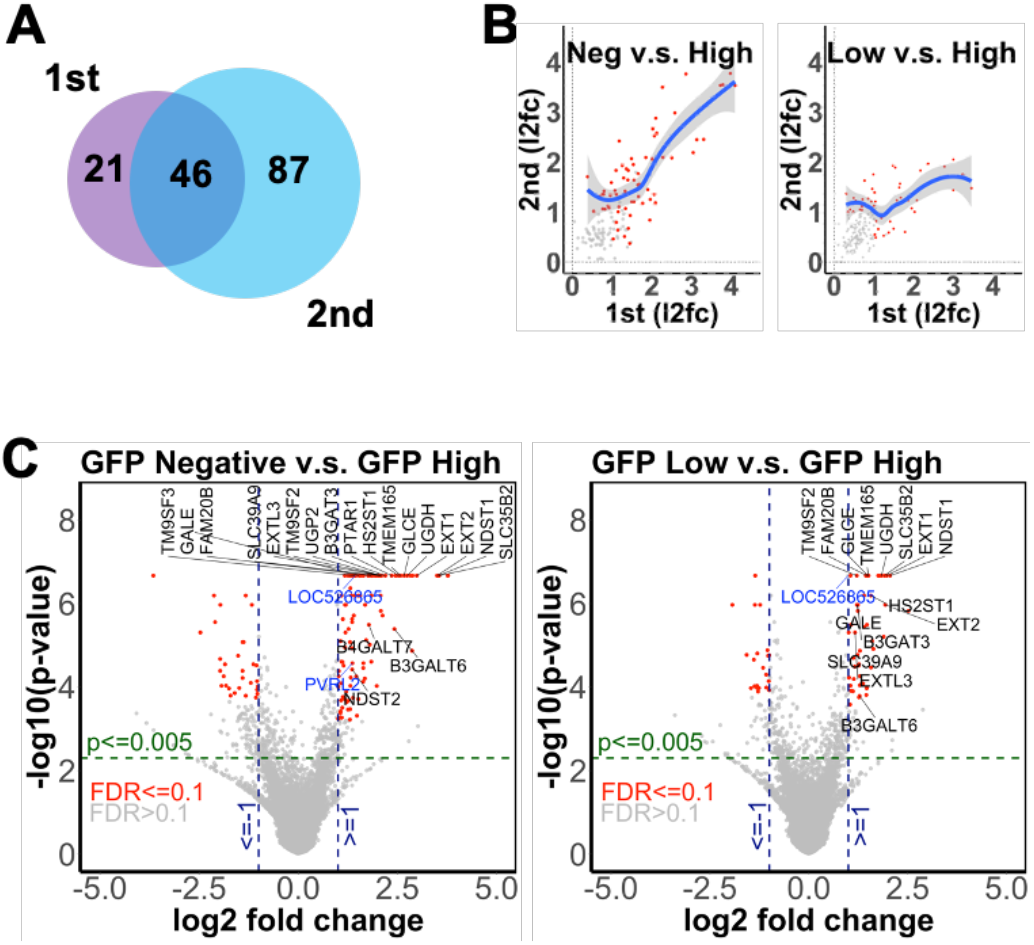
Candidate pro-viral host genes identified from the CRISPRko screens. **A.** Venn diagram showing numbers of unique and common candidates identified from the 1^st^ and 2^nd^ screen (**Data file 4**). **B.** l2fc of guides targeting genes in GFP Negative cells(left) or Low cells(right) compared to GFP Highs from the 1^st^ screen plotted against results from the 2^nd^ screen (**Data file 2,3**). All dots represent genes with p-value <=0.05 from both screens, dots highlighted in red are candidates with p-value<=0.005, FDR <0.1 and l2fc>=1 for both screens. The trendlines were drawn using Local Polynomial Regression Fitting with 0.95 as level of confidence interval. **C.** l2fc of guides for all genes in the GFP Negative(left) or Low (right) cells compared to High from the 2nd screen. Red dots represent candidates with p<0.005, FDR<0.1 and abs(l2fc)>=1 as statistical cut-offs (**Data file 3**). Genes with blue labels encode cell surface proteins previously reported as receptors for viral entry, those with black labels are candidates involved in HS biosynthesis. Blue vertical lines delineate l2fc>=1 or l2fc<=−1 boundaries.

For both screens, we compared CRISPR copy numbers from GFP Negative and Low cells to those from the GFP High cells. With strict cut-offs at adjusted p-value <= 0.005, FDR<= 0.1 and log2 Fold Changes(l2fc) >= 0.95, a total of 154 pro-viral candidate genes emerged (**Data file 3,4**, highlighted in blue). Guides targeting these genes were enriched in the GFP Negative and Low cells, with up to 25-fold increase in abundance (l2fc=4.66), indicating significant depletion of target genes. The protective effects from the gene depletion resulting in reduced GFP signal suggests pro-viral role for these candidates. The list combines 50 and 45 candidates identified from the Negative v.s. High group and the Low versus High group of the 1^st^ screen, as well as 116 and 53 (**Data file 4**) from the same groups of the 2^nd^ screen (**Fig. 4C**, highlighted in red in left and right plots; **Data file 4**). The 2^nd^ screen re-identified 46 out of the 67 pro-viral candidates from the 1^st^ screen, together with 87 new candidates (**Fig. 4A, Data file 4**). When we compared the l2fc of common pro-viral candidates from both screens, there was a good regression trend between the screens (**Fig. 4B, Data file 3,4**), especially for those identified from the GFP Negative cells (left plot), an indication of screen reproducibility and candidate reliability.

The screens identified many genes with proven or novel roles in BHV-1 cell entry. We recovered PVR (LOC526865 or nectin-1) and PVRL2 (nectin-2, **Fig. 4C**, blue labels, **Data file 4**), two membrane proteins previously implicated as receptors for BHV-1^8–10^. The screens also captured at least 20 genes involved in crucial steps of Heparan Sulfation^50^ (**Fig. 4C**, black labels, **Fig. S13,S14**). Another group of candidates, COG1, COG2, COG4, COG5, COG6 and COG7, codes for six of the eight subunits of the Conserved Oligomeric Golgi Complex (COG), an evolutionarily conserved peripheral membrane protein complex residing within the Golgi apparatus and thought to act as a retrograde vesicle tethering factor in intra-Golgi trafficking^51^. It is believed to be crucial for glycoprotein modification and Heparan Sulfate secretion to the cell surface^52^. We subsequently validated some of these candidates but due to space constraint, we will publish the detailed validation results and further studies in a separate manuscript. The identification of many such known genes and multiple subunits from the same complexes or pathways demonstrate robustness of our screen setup.

Upon release into the cytoplasm, the capsid travels to the nuclear peripheral and docks at the nuclear pore complex (NPC) for genome delivery. The NPC is composed of 34 distinct nucleoporins embedded in the NM and is a gateway for bi-directional and selective transport of both nucleic acids and peptides. From the 2nd screen, guides targeting three of the nucleoporins, NUP35, NUP50 and SEHL1 were significantly enriched in the GFP Negative cells (**Fig. 5, Data file 4**). Another hit, KPNA2 encodes for importin α1 and has been shown to mediate import of several HSV-1 proteins into the nucleus and modulate assembly and egress of its capsids^53^. XPO6, a gene encoding Exportin-6 that acquires cargo in the nucleus and shuttles it to the cytoplasm via the NPC, was also discovered (in 1st screen). How these proteins participate in the bi-directional transport of BHV-1 transcripts and proteins in and out of the nucleus remains to be studied.

**Figure 5.**
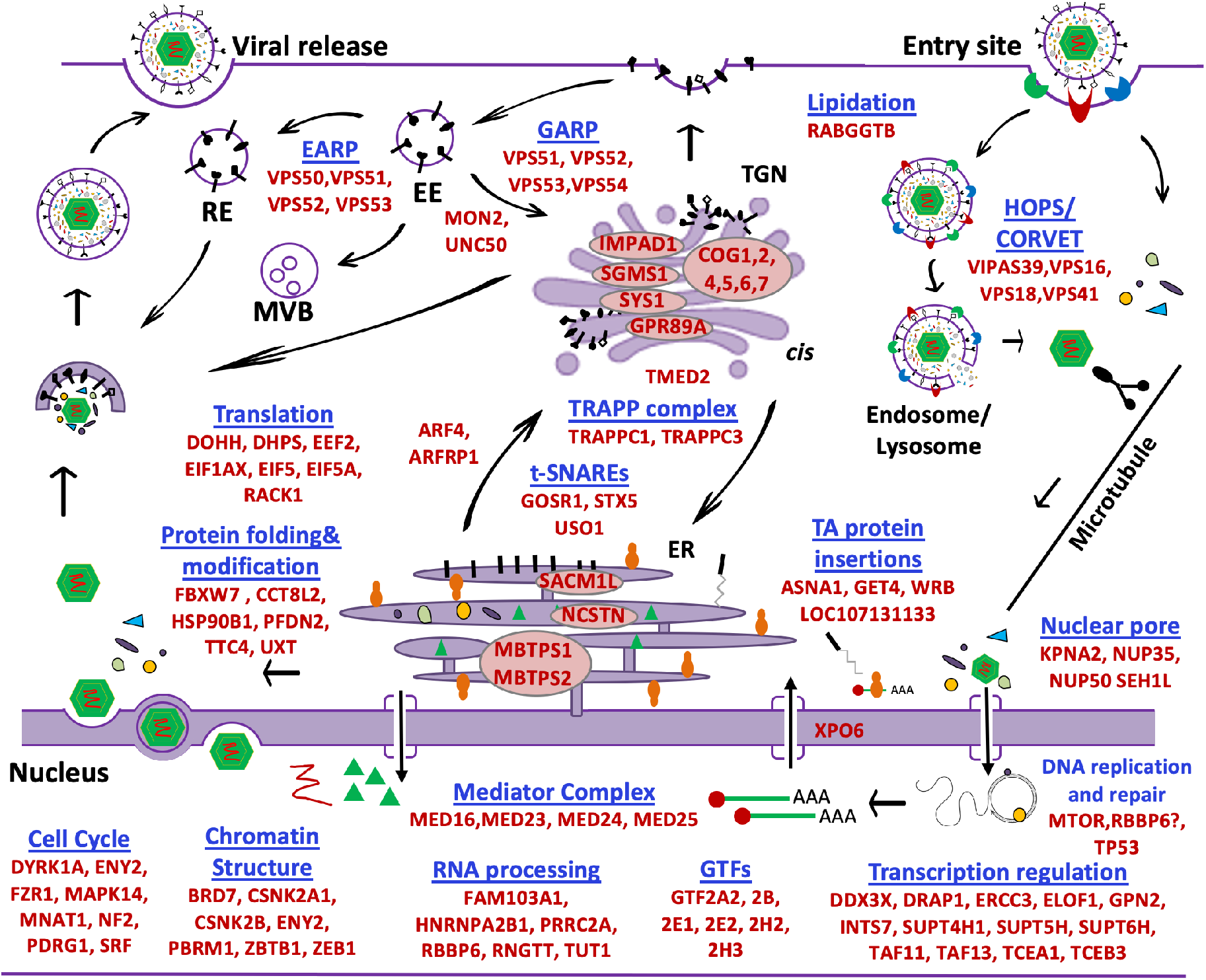
Diagram highlighting some of the candidate pro- and anti-viral host genes involved in multiple steps of the virus replication cycle. Not all candidates identified using p <0.005, FDR <=0.1 and |l2fc| >=1 as selection criteria are shown (**Data file 4**). Diagram depicts the complete life cycle of the virus from cellular entry to genome replication in the nucleus, capid egress through the double NM and cytoplasmic packaging of the capsid and teguments with host derived membrane prior to exocytosis.

Alpha herpesviruses have evolved to usurp host transcription machinery to express their own genes and manipulate host gene transcription. At least eight factors within the pre-initiation complex (PIC) were significantly depleted in the GFP Negative and Low cells from our screen (**Fig. 5**, **Data file 4**). In addition, MED16, MED23, MED24 and MED25 encoding four subunits of the Mediator complex known to interact with IE gene product ICP4 of HSV1 during active DNA replication and transcription^54^, were among some of the most depleted genes, with up to 17-fold increase in gRNA copy number (**Fig. 5**, **Data file 4**). Genes known to regulate RNA elongation and processing after transcription initiation, e.g. DDX3X^55^, DRAP1, ERCC3, GPN2, INTS7, SUPT4H1, SUPT5H, SUPT6H^54^, TCEA1 and TCEB2, were also significantly deleted. Furthermore, multiple genes involved in chromatin restructuring, namely BRD7^56^, CSNK2A1, CSNK2B, ENY2, PBRM1^56^, ZBTB1, and ZEB1, were also identified. Many of these gene products have been found to influence HSV-1 replication and transcription^54^; BHV-1 likely hijacks them for viral gene expression in a similar fashion as HSV-1.

During and post transcription, precursor mRNA (pre-mRNA) transcripts are capped, poly-Adenylated, and spliced prior to nuclear export. Many animal viruses hijack host factors to achieve these modifications for efficient viral translation or divert host RNA for degradation via virion host shutoff (vhs)^57^. Indeed, several genes including FAM103A1 and RNGTT important for mRNA capping^58^, HNRPA2B1^59^ and PRRC2A^60^ for m6A-dependent nuclear RNA processing, and RBBP6 for pre-mRNA processing were all preferentially deleted in GFP Negative and Low cells (**Fig. 5, Data file 4**). In addition to factors directly involved in mRNA processing, TUT1, a gene encoding a protein that adds 3’ polyA tails to mRNA and polyU tails to miRNAs and snRNAs, the latter being important for splicing of pre-mRNAs^61^, was also depleted.

Once transcribed and modified, most viral transcripts are exported to the cytoplasm to be translated into peptides that are folded and transported prior to viral packaging and egress. Protein translation is a multi-step process consists of initiation, chain elongation and termination; our screen identified multiple factors namely EEF2, EIF1AX, EIF5, EIF5A involved in these processes. Interestingly, genes encoding two proteins catalysing the conversion of lysine to the unique amino acid hypusine in EIF5A^62^, DOHH and DHPS (**Fig. 5, Data file 4**), were significantly depleted in GFP Negative and Low cells from both screens, potentially highlighting the prime importance of EIF5A in modulating BHV-1 protein translation. In addition to translation initiation and control, genes encoding factors important for proper folding of newly synthesized peptides, e.g. TTC4, HSP90B1, PFDN2, were also preferentially deleted in GFP Negative and Low cells.

Another interesting cohort of candidates encodes for factors that promote intracellular protein and membrane trafficking. In addition to the COG complex, two other clusters of genes were enriched from our screens based on STRING protein analysis^63^. One cluster consists of VPS51, VPS52, and VPS53 (**Fig. 6C**, red nodes), genes encoding subunits of the Golgi-associated retrograde protein (GARP), a retromer complex that tethers retrograde endosomal carriers to the trans-Golgi network (TGN)^64^. A few genes known to regulate this retrograde trafficking process, namely SYS1, ARFRP1, and RGP1, were also enriched. The fourth subunit of GARP, VPS54 locates the complex to the TGN and to recycling endosomes to a lesser extent^64^. Replacing VPS54 with VPS50 forms another complex, endosome-associated recycling protein or EARP that primarily resides on recycling endosomes^64^. The other cluster of genes code for tethering or regulatory factors essential for intra-Golgi transport or membrane trafficking between the Endoplasmic Reticulum (ER) and the Golgi Apparatus (**Fig. 6C**, green nodes). These include members of tethering SNAP Receptors or t-SNAREs (GOSR1 and STX5), regulator of SNARE assembly required for ER-to-Golgi transport (USO1 and ARF4) and subunits of the transport protein particle tethering complex or the TRAPP complex (TRAPPC1 and TRAPPC3). There are also nodes disconnected from the network with suspected roles in intra-cellular trafficking, namely GPR89A, RABGGTB, SGMS1, UNC50, and VPS13D (**Fig. 6C**, blue nodes, **Data file 4**).

**Figure 6.**
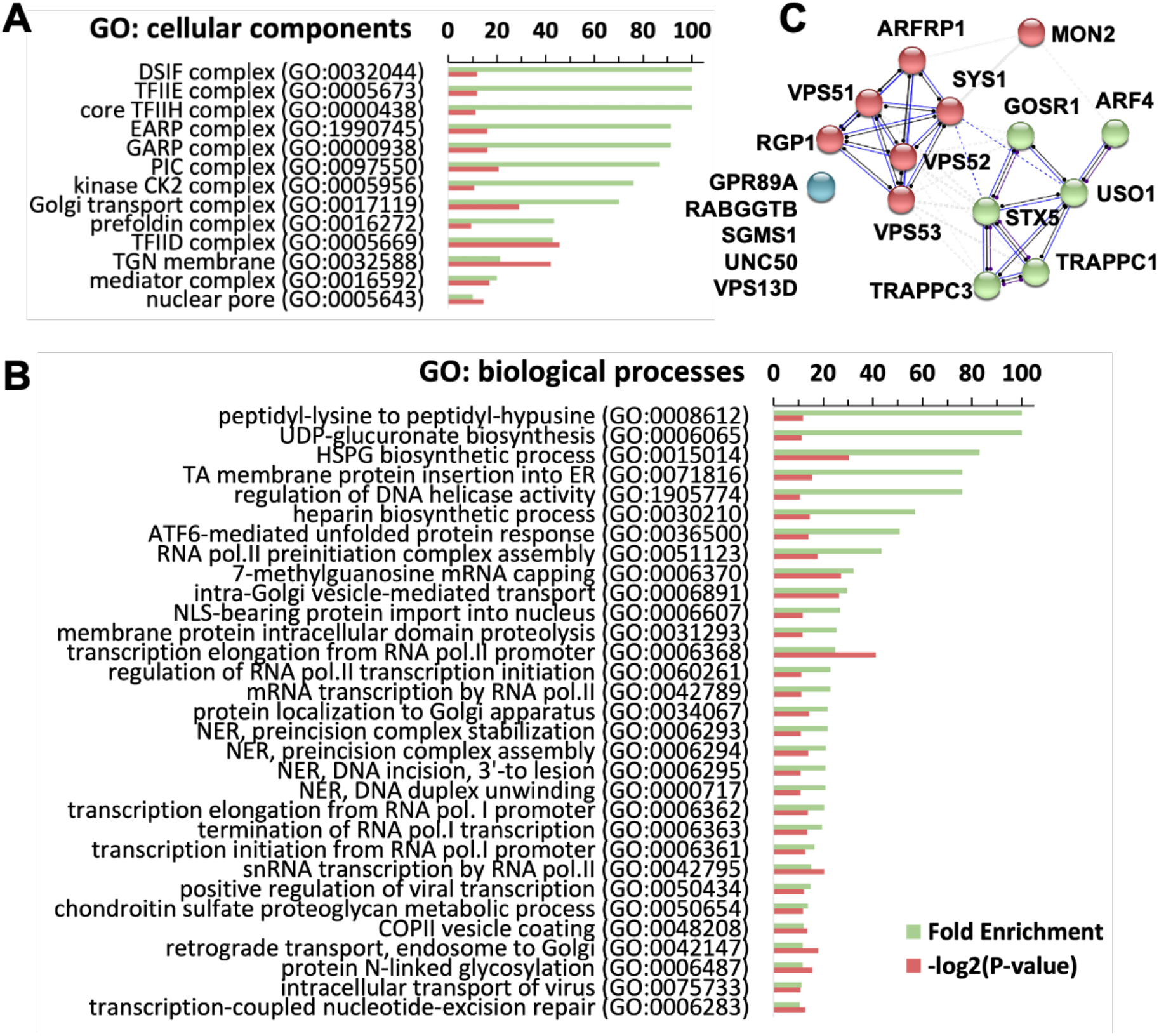
Analyses of enriched cell components and biological processes using GO or STRING. **A.** Gene Ontology analysis (GO) reveals the top enriched cellular components candidate genes involved in virus replication. Cellular components shown have Fold Enrichment >=10, and −log2(p-value)>=10 (To see full list, please refer to **Data file 5**). **B.** GO reveals the top biological processes that can be important for the virus using the same statistical cut-off as in A (To see full list, please refer to **Data file 5**). **C.** STRING analysis and clustering of some of the candidates involved in intra-cellular trafficking. Blue dots represent disconnected genes based on the current STRING database 11.0.

### CRISPR knockout screen also identifies anti-viral candidates

Using similar statistical cut-offs (p<0.005, FDR<=0.1, l2fc<=−0.95) as used to filter for pro-viral candidates, we identified 56 anti-viral candidate genes from both screens. Initial analysis and literature search highlighted several candidates of interest e.g. CCND1, CDC37, FBXW11 and TFDP1 that regulate cell cycle progression, CDC45, MCM2 and MCM3 as members of the minichromosome maintenance proteins, Furin that cleaves selected glycoproteins, TYMS that encodes for Thymidylate Synthetase, and UBE2M for NEDD8-conjugating enzyme UBC12 (**Data file 4**). By slightly relaxing the statistics (p<=0.05, FDR<=0.3 and l2fc <= −0.75), we extended the list to include 86 more candidates (**Data file 4**). It expands the group of genes that regulate cell cycle and DNA replication and highlights other sets of genes with good anti-viral potential. To name a few, genes associated with the HOPS/CORVETT complexes that coordinate late endosome and lysosome fusion, VIPAS39, VPS16, VPS18, and VPS41, became significant (**Fig. 5**). In addition, the screen also pinpointed other E2 ubiquitin-conjugating enzymes or ubiquitin related or like proteins, including UBA52, UBL5, UBE2D3, with the latter shown to be essential for RIG-I and MAVS aggregation in antiviral innate immunity^65^. Two negative regulators of mTORC1 signalling, TSC1 and TSC2 that are phosphorylated by Us3 during HSV-1 infection resulting in mTORC1 activation^66,67^, were also identified.

### Gene Ontology analyses identify enriched pro-viral cell components and biological processes

To prioritize validation efforts, we conducted formal gene ontology (GO) analyses to help determine which cellular components and biological processes are the most enriched. With Fold Enrichment >10 and −log2(P-value) >10 as cut-offs, 13 cellular components (**Fig. 6A**) and 30 biological processes (**Fig. 6B**) stood out from the list of pro-viral candidates (**Data files 4,5**). Both lists contain components or processes with diverse biological functions occurring at different locations in the host cell. Some of these candidates are obvious and already highlighted above, such as processes at different steps of HS synthesis (e.g. GO:0006065, GO:0015014, and GO:0030210) and general transcription factors indispensable for viral transcription (e.g. GO:0005673, GO:0000438, GO:0051123, GO:0006368), while others require substantial work to pinpoint their possible function in BHV-1 biology. Two components captured our prime attention, the EARP complex and the GARP complex (GO:1990745 and GO:0000938) being the 4^th^ and 5^th^ most enriched cellular components according to GO (**Fig. 6A**). Both of them have been shown to be important for endocytic recycling^64^ but no roles in alpha-herpesvirus replication has been documented, and we decided to explore how and at what stages of the BHV-1 life cycle they are recruited by the virus.

### VPS51, VPS52, and VPS53 knockout severely impairs production of infectious particles

Based on the screens the three subunits shared by GARP and EARP, i.e. VPS51, VPS52 and VPS53, were preferentially depleted in the GFP Negative and Low samples, with targeting guides enriched by up to 4-fold (**Data file 4**). To validate these results, we produced individual KO MDBK cell lines lacking VPS51, VPS52 or VPS53 (**Table S5**) and performed plaque assays on them. The cells became substantially less permissive to BHV-1 infection, with virus titre dropping by at least 274-fold (VPS52KO) to almost undetectable (VPS51KO, VPS53KO) (**Fig. 7A**). The plaques also became much smaller, with at least 80% reduction in size compared to those grown in control cells (WT, **Fig. 7B**). Overexpressing cDNA encoding the missing subunits in corresponding KO cells (VPS51R, VPS52R, VPS53R) completed remedied the impairment, in both viral titre (**Fig. 7A**) and plaque size (**Fig. 7B**). By measuring GFP fluorescent intensity from 0 h.p.i. to 72 h.p.i, we also tracked the rate of VP26 synthesis in the KO and rescue cells infected with the tagged virus at MOI =3 (**Fig. 7C**) or MOI =0.1 (**Fig. 7E**). While the GFP growth rates in the rescue (VPS52R) were almost identical to that in the control cells (Cas9+/+), it was reduced to 1/3 in the KO cells in the MOI =3 group after just two cycles of infection, measured between 16 h.p.i. and 24 h.p.i (**Fig. 7C**). The reduction was even more profound in the MOI =0.1 group, as GFP signal barely increased throughout the infection in the KO cells compared to the rescue (VPS52R) and the Cas9+/+ control (**Fig. 7E**). The plaque assay results were replicated by infecting these cells with the gamma-herpesvirus Alcelaphine Herpesvirus 1 (**Fig. S15**)^68^.

**Figure 7.**
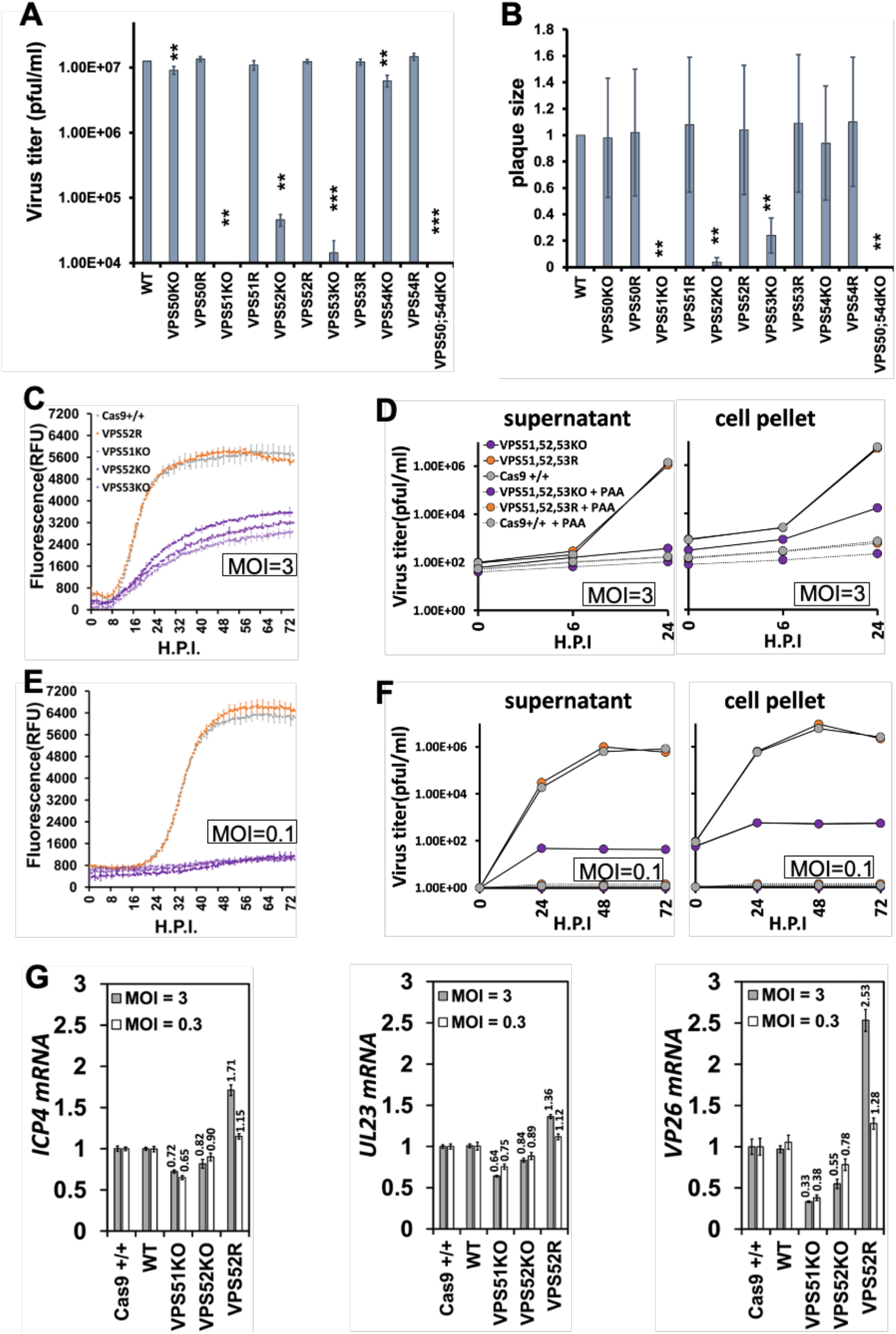
The loss of both EARP and GARP impairs virus replication and the two complexes compensate each other. **A,B.** Plaque assay results using cells with single gene KOs (VPS50KO to VPS54KO), KO rescues (VPS50R to VPS54R), double KOs (VPS50;54dKO) or wild type cells(wt). Data presented are averages of three repeats (n=3) from at least three clones for each KO. **: p<0.05; ***: p<0.005. **C,E.** VP26-GFP growth curves based on fluorescence in single gene KO cells (VPS51KO to VPS53KO), rescue cells (VPS52R) or Cas9+/+ cells infected with the virus at MOI=3 or MOI=0.1 Data were collected from at least three independent KO clones (n=3) for each gene. **D,F.** Plaque assay results using supernatants or cell pellets collected from single gene KO cells (VPS51,52,53KO), KO rescues (VPS51,52,53R) or Cas9+/+ cells infected with BHV-1 at MOI=3 or 0.1, with or without treatment of PAA, a drug that blocks viral genome replication and progression of virus infection cycle. VPS51,52,53KO and VPS51,52,53R represent averages from all KO samples or rescues; the error bars are smaller than the markers for most data points thus are not visible. **G.** relative expression levels of immediate early (ICP4), early (UL23) and late (VP26) viral transcipts from Cas9+/+, wild type (WT), KO (VPS51KO, VPS52KO) or rescue (VPS52R) cells at 6 h.p.i. with BHV-1 at MOI=3(grey) or 0.3(white). Data were the results of three technical repeats of samples from a single experiment.

To further dissect the phenomenon of reduced virus production, we quantified cell associated and cell free virus produced from these KO cells by titrating supernatant and cell pellet samples collected at multiple time points on WT MDBKs. Throughout a 24-hour infection with BHV-1 at MOI =3, the KO cells (VPS51,52,53KO) almost completed halted the production of cell free virus, with nominal increase (6.3-fold) in plaque forming units (PFUs) in the supernatants compared to that from the Cas9+/+ cells (14737-fold) and the rescues (11167-fold, VPS51,52,53R), this 2339-fold reduction (relative to Cas9+/+) was comparable to that imposed by PAA blockage of viral genome replication (**Fig. 7D**, left). At 0h.p.i, the amount of cell associated PFUs in the KO cells decreased by 3.2-fold compared to Cas9+/+ after infection at MOI =3 (p<0.0001) and by 1.5-fold at MOI =0.1 (p =0.029) (**Fig. 7D**, right). The subsequent production of cell associated infectious virus particles was also greatly affected with a 123-fold decrease by 24hpi (**Fig. 7D**, right), 19 times less pronounced than the 2339-fold drop in the supernatant as the KO cells still produced virus in the pellets with a 58-fold increase over 0 h.p.i (**Fig. 7D**, right). The trend was similar for cells infected at MOI =0.1; by 24 h.p.i, the amount of cell free PFUs produced from the KO cells was 389-fold less than that from Cas9+/+ (**Fig. 7F**, left) and cell associated PFUs was 666-fold less (**Fig. 7F**, right); and beyond 24 h.p.i the KO cells barely produced any more infectious virus particles. Taken together, these initial data confirm that the KO of VPS51, VPS52, and VPS53 greatly impairs BHV-1 replication in MDBK cells. The loss of both GARP and EARP leads to reduced production of both cell free and cell associated infectious BHV-1 particles, with cell free virus more severely impacted by the KO.

### VPS50 and VPS54 double KO replicates the effects of VPS51, VPS52 and VPS53 KO

Because GARP and EARP share these three subunits, the KO of any of them leads to loss of both complexes, and our data have shown that at least one of them is important for efficient alpha- and gamma-herpesvirus replication. To investigate this, we first created EARP or GARP specific KO cell lines by deleting just VPS50 (VPS50KO) or VPS54 (VPS54KO) (**Table S5**). Interestingly, when we conducted plaque assays on them there was only a small reduction (1.4-fold in VPS50KO, p=0.015; and 2-fold in VPS54KO, p=0.014) in virus titre compared to the WT (**Fig. 7A**), and no change in plaque sizes was observed for both series of KOs (**Fig. 7B**). Although still significant, the decrease in titre of virus grown in VPS50KO or VPS54KO was very modest compared to that observed in VPS51KO, VPS52KO and VPS53KO. These results suggest that both GARP and EARP can be recruited by the virus, possibly via independent routes for similar purposes or they are functionally redundant through the same pathway. To test this, we produced VPS50;54dKO cell clones missing both VPS50 and VPS54 (**Table S5**) and conducted plaque assays as before in parallel to the single KO clones. The dKO cell clones recapitulated the results obtained from VPS51KO, VPS52KO and VPS53KO clones perfectly with no plaques formed (**Fig. 7A, B**). These experiments suggest that the roles GARP and EARP play relative to the BHV-1 virus biology are almost interchangeable and at least one of them is required for efficient virus replication.

### VPS51 KO leads to reduction in viral genome and viral transcript copy numbers

To help pinpoint the cause for impaired virus production, we conducted a pilot experiment to see if cell entry or viral gDNA replication could be affected in KO cells missing both GARP and EARP. We quantified viral genomic DNA (gDNA) using reverse transcription and qPCR (RT-qPCR) and then compared total viral gDNA levels in cell pellets collected from KO cells (VPS51KO and VPS52KO), control cells (WT and Cas9+/+) or rescues (VPS52R) at 6 h.p.i (**Fig. S16**). We found that there was reduction of viral gDNA levels in KO cells with a 2-fold drop for VPS51KO (p = 0.013) and 1.2-fold for VPS52KO (p = 0.043) at MOI=3; and 1.5-fold (p=0.0001) and 1.1-fold (p = 0.00013) drop at MOI=0.3 respectively compared to Cas9+/+. Interestingly, total viral gDNA levels in the rescue cells (VPS52R) with cDNA overexpression increased to 2.5-fold at MOI=3 (p = 0.0002) and to 1.3-fold at MOI=0.3 (p = 0.0088) of that in the Cas9+/+ cells.

In addition to viral gDNA, we also investigated whether the loss of GARP and EARP negatively impacted transcripts from the Immediate Early (IE), Early (E) and Late (L) viral genes which in turn might have caused the dramatic reduction of viral yield. Using RT-qPCR, we quantified mRNA transcribed from ICP4 (IE), UL23 (E), and VP26 (L) in cell pellets harvested from GARP/EARP KO cells (VPS51KO, VPS52KO), control cells (Cas9+/+, WT) and rescue cells (VPS52R) at 6 h.p.i (**Fig. 7G**). For both infections with BHV-1 at MOI = 3 or MOI = 0.1, we observed reduced mRNA levels of ICP4, UL23, and VP26, ranging from 25% (1.3-fold for UL23 at MOI=0.3, p=0.0006) to 67% (3-fold, for VP26 at MOI=3, p=0.0003) decrease in VPS51KO cells compared to those in Cas9+/+, with severity steadily increasing through the three stages of viral gene transcription. The same trend was observed in the VPS52KO cells, but it was reversed in the VPS52R rescue cells. The higher abundance of viral gDNA in VPS52R (**Fig. S16**) was coupled with enhanced mRNA levels, with up to 2.5-fold increase (VP26 at MOI=3, p=0.0001) for all the three genes tested (**Fig. 7G**).

### VPS51 KO potentially causes VP26 retention and the formation of larger but fewer capsid aggregates in the nucleus

After showing that the double loss of GARP and EARP restricted the virus on the DNA and mRNA level, we proceeded to examine whether and how viral proteins could also be affected. Due to the lack of BHV-1 specific primary antibodies, we chose to monitor the dynamics of the GFP-tagged VP26, the small capsid surface protein that translocates into the nucleus for capsid assembly and genome packaging. By tracing GFP, we examined the status of VP26 hoping to gain insight into the viral capsid prior to and during nuclear egress at the late stage of infection. To achieve this, we first infected VPS51KO and Cas9+/+ cells with the GFP tagged BHV-1 virus at a MOI = 0.3 and visualized VP26-GFP in relation to the nuclei by confocal microscopy at 6 h.p.i. We found that compared to Cas9+/+ control cells, the majority of VP26 in VPS51KO cells was confined to the nuclei (**Fig. 8**). In addition, the VP26 protein formed fewer and more sparsed aggregates in the VPS51KO cells; however the aggregates in these cells are much bigger in size compared to the controls. These preliminary results seem to suggest that the loss of GARP and EARP lead to abnormaties in the dynamics of the VP26 protein and probably that of capsid assembly prior to nuclear egress.

**Figure 8.**
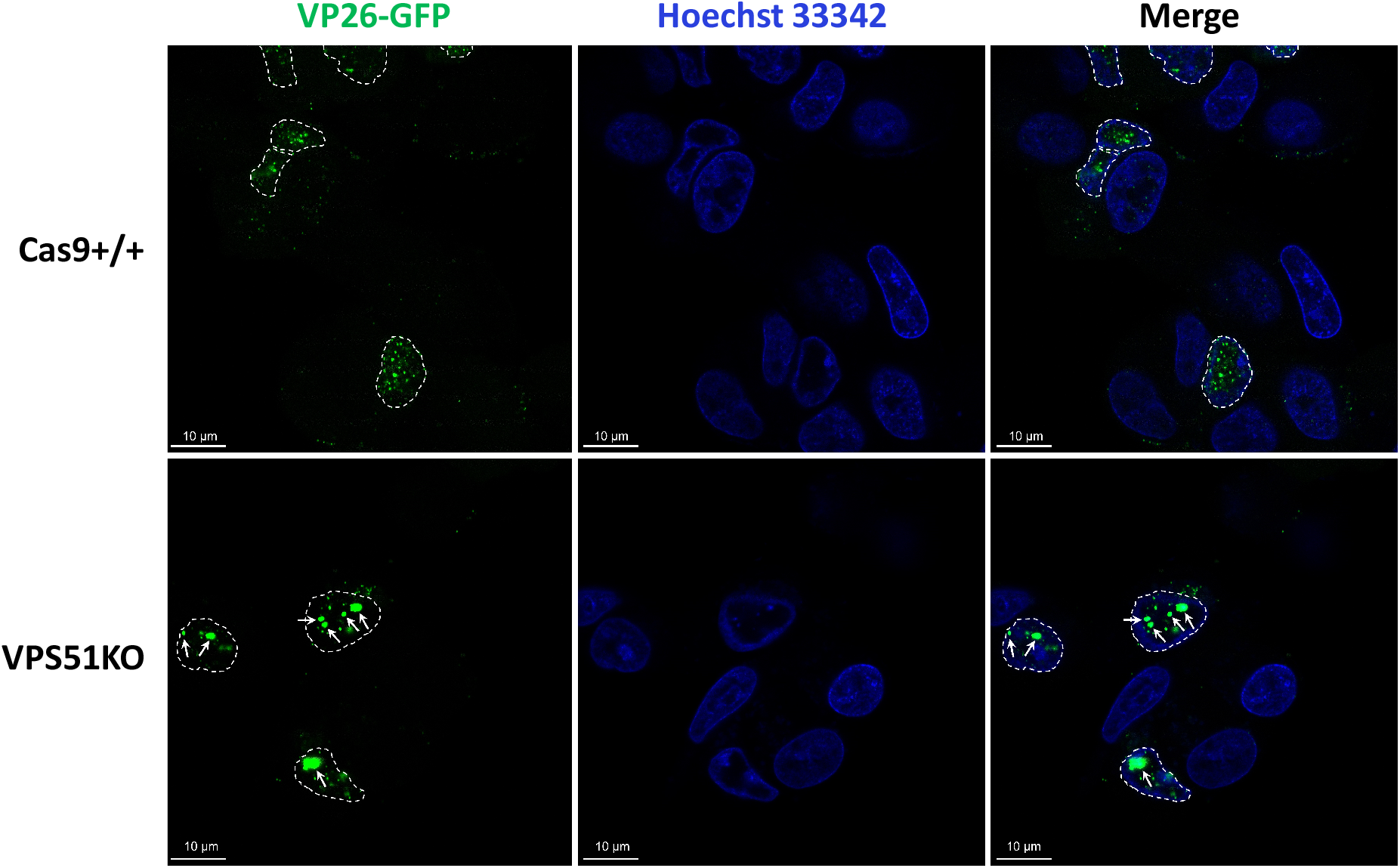
The loss of EARP and GARP leads to retention of capsids in the nuclei and formation of larger but fewer aggregates. Retention of VP26-GFP as large aggregates (white arrows) in the nuclei in VPS51KO (left) cells compared to small punctates in Cas9+/+ cells (right). At 6 h.p.i with the VP26 tagged BHV-1 virus (MOI= 0.3), the localization of VP26-GFP was captured by Zeiss LSM880 confocal microscope while the nuclei were identified with Hoechst 33342. Original magnification: ×60; Bars = 10 μm.

## Discussion

By conducting a genome wide CRISPR knockout screen using the very effective library created in this study, we identified many interesting cow genes that could affect various aspects of the BHV-1 life cycle. Some of the most intriguing examples encode for subunits of the EARP and GARP complexes known for promoting endosome recycling^64^. Prior to this publication there had been no documentation on the involvement of these complexes in alpha-herpesvirus replication. After validating the screening results, we conducted a series of experiments to identify potential mechanisms and routes these complexes are recruited to the benefit of the virus. The evidence we have gathered points in three general directions: cell entry or capsid trafficking to the nucleus, nuclear egress and secondary envelopment.

Our data suggest that the loss of EARP and GARP affects virus entry or transport to the nucleus. By infecting KO cells missing both EARP and GARP, we observed up to 3-fold reduction of viral transcripts at 6h.p.i compared to the control cells (**Fig. 7G**). Since ICP4 and UL23 transcripts are primarily produced from the original viral gDNA prior to genome replication, their reduction is likely caused by the decrease of viral gDNA reaching the nucleus, assuming that viral transcription efficiency is not altered at these loci. Conversely, when the VPS52 subunit was overexpressed, the total viral genome quantity increased instead (**Fig. S16**), along with the IE and E transcripts from all three stages of viral gene expression (**Fig. 7G**). This is perhaps best explained by enhanced cell entry or transport to the nucleus possibly caused by increased GARP/EARP abundance in the cytoplasm due to VPS52 overexpression, further strengthening this proposition. Lastly, we also observed a higher magnitude of changes of VP26 mRNA levels in VPS52KO or VPS52R than those of ICP4 and UL23 mRNA (**Fig. 7G**). As a major viral transcription regulator, mild changes in ICP4 could lead to more dramatic changes of downstream genes including UL23 and VP26. We will repeat both the viral gDNA and transcript quantification experiments in the future to see if the results described above are reproducible.

Despite the likelihood that cytoplasmic entry and or capsid trafficking to the nucleus is affected by the KO, this alone could only explain a small portion of the virus titre loss. The absence of GARP and EARP lead to at least a 274-fold decrease of total infectious virus and 80% reduction in plaque size (**Fig. 7A, B**), but only up to 3-fold can potentially be attributed to defects in cell or nuclear entry according to the viral gDNA, ICP4 mRNA and UL26 mRNA copy number changes at 6h.p.i (**Fig. S16, 7G**). Throughout the course of a 24-hour infection at MOI=3, VPS51KO cells barely released any infectious new virus into the supernatant (6.3-fold increase, **Fig. 7D**), while the titre of virus from Cas9+/+ increased exponentially (14737-fold), a 2339-fold difference between the two. The production of cell associated virus has also been affected by the KO, albeit not as severely. The titre of cell associated virus released by freeze and thaw from pellets of infected KO cells did rise by 58-fold, although it was still substantially lower than the 7127-fold increase in Cas9+/+ control cells. Taken together, these data show that the majority of infectious new virions produced in VPS51KO, VPS52KO, and VPS53KO cells were cell associated, and at a remarkably reduced level. It could be that the majority of cell free virus particles released by the KO cells were non-infectious due to unknown defects.

Vesicles of TGN origin^69^ and recycling endosomes have both been strongly indicated as assembly sites for cytoplasmic envelopment of BHV-1^25^, although the involvement of TGN is heavily debated for HSV-1. Alpha-herpesvirus genomes encode a dozen glycoproteins and many of them are secreted and concentrated to the PM and recycled back by endocytosis; for HSV-1, they include the essential glycoproteins i.e. gB, gD, gH/L and others^21,70^. These glycoproteins are packaged into virions during secondary envelopment and are essential for successful cell entry by mediating membrane fusion^71^. It has been shown that some viral glycoproteins such as gH of HSV-1 localize to vesicles positive for TGN46, a TGN marker, or on recycling endosomes^20,21^. Schindler et al showed that GARP mainly resides on the TGN and EARP at recycling endosomes^64^. Co-localization of the GARP and EARP with the glycoproteins suggests that these complexes mediate capsid and tegument wrapping with these vesicles decorated with gB, gH/L and others essential for cell entry. New cell free virions produced from the KO cells lack such glycoproteins due to the KO inflicted absence of fusion between EE derived microtubules and RE or TGN prior to wrapping, disabling them for a second-round infection. Schindler et al also found that GARP mainly resides on TGN but can also be present on REs, potentially mediating fusion of EE derived vesicles with both RE and the TGN. Knocking out TGN specific tethering complexes in cells lacking VPS50 and EARP, such as VAMP4^72^ can help determine whether one or both routes mediated by GARP are used by the virus; the co-discovery of candidate genes involved in EE to TGN trafficking such as ARFRP1, RGP1 and SYS1 seems to support the latter.

Based on our own data and existing evidence, we developed a model for BHV-1 secondary envelopment. Under normal condition (**Fig. 9**, shaded grey), viral glycoproteins are endocytosed from the cell membrane, membrane microtubules carrying these glycoproteins bud from the EEs and fuse to the TGN via GARP mediated retrograde transport, to REs in the fast recycling route tethered by EARP and or GARP or to REs in the slow recycling route regulated by Rab11. Membrane vesicles budded from the TGNs or those that result from fusions to REs are then used by the virus to wrap capsids and tegument proteins into virions for exocytosis through natural endosomal recycling. Alternatively, EEs can become multi-vesicle bodies (MVBs) and then late endosomes (LE) for protein degradation. In addition, some Rab11 positive slow REs^73^ can also morph into initial autophagic vacuoles (AVi)^73^ instead of being recycled back to the PM. In cells without functional GARP and EARP tethering complexes however, (**Fig. 9**, shaded blue), microtubules derived from EEs are unable to fuse with the TGN or REs bound for fast recycling, depriving the virus access to glycoprotein containing fast recycling REs or TGN derived vesicles for secondary envelopment. Instead, the virus primarily uses membrane vesicles deficient of viral glycoproteins for envelopment, producing defective virions without glycoproteins important for cell entry, thus these virions are incapable of starting a new cycle of infection. Due to the lack of GARP and EARP, EEs carrying these glycoproteins are more likely to become LEs or AVis for degradation. The assembly and wrapping of cell associated viruses in the cytoplasm might use a different mechanism, thus not affected by the KO and can still produce infectious virions. Although this model is developed for BHV-1, it could also be applicable to other herpesviruses such as Alcelaphine herpesvirus as suggested by our data (**Fig. S15**).

**Figure 9.**
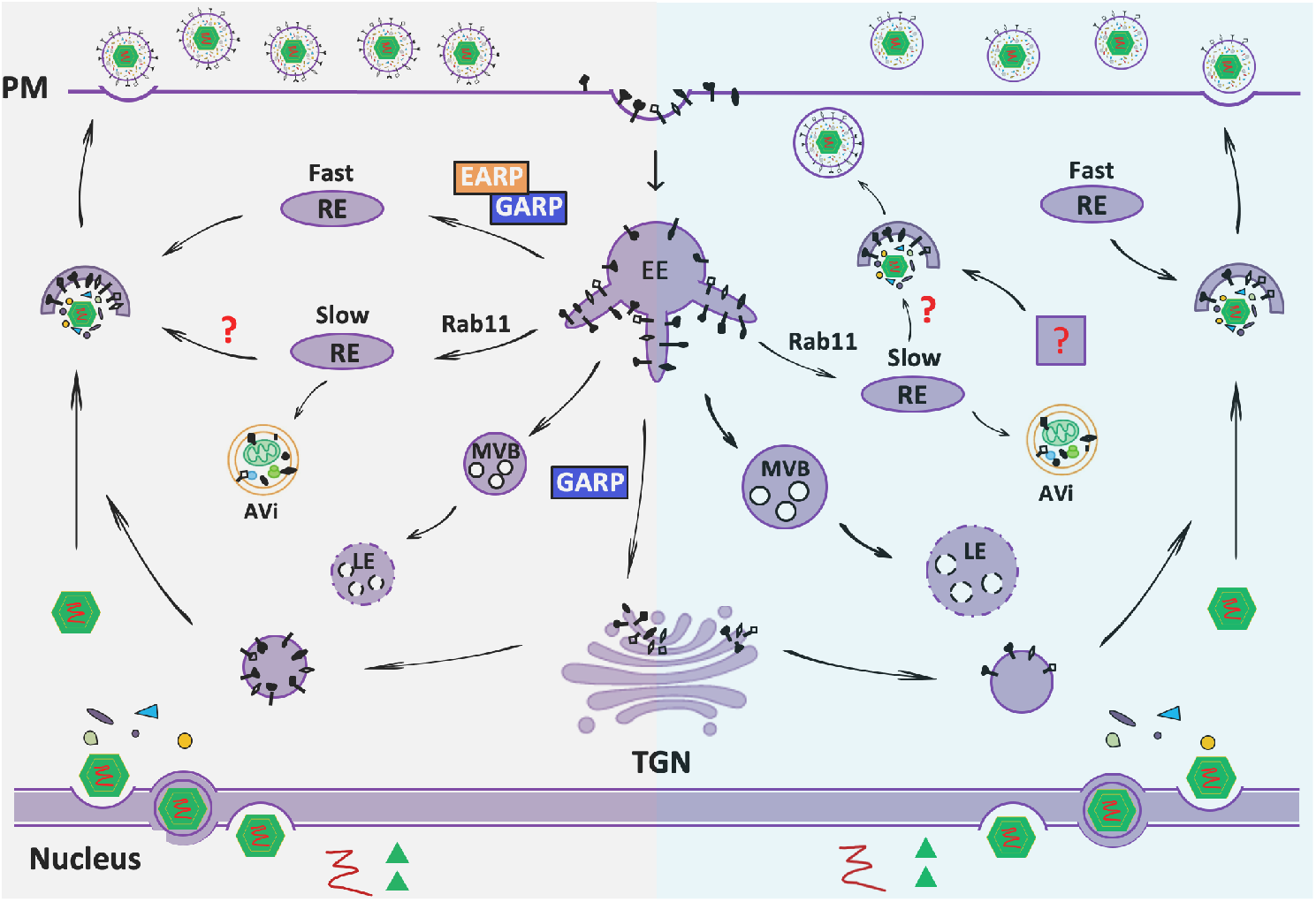
Predicted model for secondary envelopment of capsids in the cytoplasm during BHV-1 infection in MDBKs. Grey shaded area represents normal condition and blue shaded area represents cell missing both EARP and GARP. AVi: initial autophagic vacuoles, EE: early endosomes; LE: late endosomes, MVB: multi-vesicle bodies, PM: plasma membrane; RE: recycling endosomes; TGN: trans-Golgi network.

Besides impairment to secondary envelopment due to loss of GARP and EARP, we also observed potential capsid retention and aggregation in the nucleus. Our preliminary data, based on a single time point at 6 h.p.i, showed that there were fewer but larger VP26 and likely capsid aggregates in the nuclei of VPS51KO cells (**Fig. 8**). This phenomenon resembles the accumulation of enveloped virions in the perinuclear space or in membrane vesicles that bulged into the nucleoplasm in cells infected with a mutant HSV-1 strain lacking both gB and gH^74^. Farnworth et al reported that as members of the core fusion machinery, not only are gB and gH important for cell entry, they are also essential for the fusion between the outer NM and the primary membrane of the capsid crossing the perinuclear space. gB and gH have also been shown to be required for nuclear egress via NM breakdown in specific cell types induced by the virus^75^. Since gB is the most highly conserved of all surface glycoproteins among alpha-herpesviruses, it is likely the combinatorial action of gB and gH is also required for nuclear exit of BHV-1 capsids. As mentioned earlier, both gB and gH are secreted to the surface of the cell, it is possible that the gB and gH that reach the outer NM at the capsid egress site are endocytosed from the cytoplasmic membrane, possibly transported via the Golgi network and ER. If true, this would explain why capsids produced in cells lacking GARP are unable to cross the outer NM via membrane fusion as the retrograde trafficking from the gB and gH carrying EE to TGN is cut off; and those that do enter the cytoplasm might achieve it by budding or physically breaking down the NM which can be less efficient processes^76^. It could also be that complementary to the model proposed above, GARP and EARP are recruited at multiple steps of the virus replication for BHV-1, and the retrograde transport from EE to TGN mediated by GARP is even more important for trafficking endocytosed glycoproteins to the NM for nuclear egress than for secondary envelopment. Although we need to examine the status of the capsids at more time points besides 6 h.p.i and the trafficking of gB and gH from cell membrane to NM is largely speculative at this stage, our screen did identify multiple candidates known for bi-direction trafficking between the Golgi and ER, such as the GOSR1, GOSR2 and USO1 of the t-SNAREs, and ARFRP1, ARL1 (**Fig. 5C**). Validating these candidates and studying their detailed mechanisms of action might help settle this speculation. Also tracking the movement of gB and gH and examining the interactions between the capsids and NMs with Electron Microscopy in GARP/EARP deficient and WT cells can offer extra clarity.

Although we focused our efforts on validating and analysing pro-viral candidate genes in this paper, anti-viral candidates can also be fascinating in terms of biology and their therapeutic potentials. The reduced sensitivity in identifying anti-viral candidates compared to the discovery of pro-viral genes could partially be due to our screen setup. Constrained by FACS sort capacity, the experiments returned a limited number of cells for each cell group, from the 1^st^ screen in particular (**Table S3,4**). It likely contributed to lower discovery rate for anti-viral candidates as they are targeted by preferentially depleted guides in the GFP Negative and Low cells. Nonetheless, our screens still recovered multiple anti-viral candidate genes that are well worth follow-up investigation, such as the many genes controlling cell cycle progression and DNA replication, subunits of the HOPS and CORVETT tethering complexes (**Fig. 5**) and the ubiquitin ligases or similar that mark proteins for degradation (**Data file 4**). One way to improve the sensitivity is to scale up, at the expense of added labour, FACS hours and downstream costs. Alternatively, a new CRISPRko library with fewer but experimentally proven guides per gene can improve screen statistics while maintaining the scale. Although not immediately attainable, the goal is achievable in the future by curating good performing guides from btCRISPRko.v1 and similar libraries in addition to improved CRISPR design algorithms and cow genome information into the new design.

The data we presented above speaks for the power of CRISPR screens. They are very effective tools for gene discovery but prior to our project, all CRISPR screens were conducted in human and model organisms^30–32^, no such library existed for livestock animals. A good understanding of host pathogen interactions in livestock is crucial for animal health, food security and the combat against spread of zoonotic diseases. The genome wide CRISPR library we built enabled us to uncover candidate genes involved in various aspects of BHV-1 replication, many of which remain to be validated and studied. We hope the gene lists produced here can be useful resource for new projects and comparative studies between herpesviruses. Like other herpes viruses, BHV-1 has co-evolved to tightly regulate the host cell niche for its own benefits, it will be important to understand the detailed mechanisms of such controls to aid development of therapeutic strategies. Due to the experimental design, we did not identify host factors important for latency, which is an important aspect of BHV-1 biology; it will be beneficial to adapt this screening strategy to gain further knowledge in this area when a suitable cell- or tissue-based latency reactivation model is developed. The screening platform we built is readily deployable to study other economically important bovine pathogens, such as the foot and mouth disease virus, bovine rotavirus, bovine viral diarrhoea virus, bovine respiratory syncytial virus or Schmallenberg virus, as long as reliable assays to isolate distinct and relevant phenotypes are established. The library will also be useful to understand other aspects of biology, such as maintenance of pluripotency, stem cell differentiation and maybe even aid efforts on repurposing BHV-1 as a cancer therapy vector^77–80^.

The library screen pipeline presented here has been very effective in host gene discovery, but there is also room for improvement. We chose MDBKs cells for the screen because a cell line more relevant to the disease manifestation does not exist; it is one of the few cell lines that support active BHV-1 infection, hence most commonly used. Although our library targets 21,156 coding genes, in reality only those actively expressed or induced by BHV-1 in the MDBKs were screened in this study, and to look into those inadvertently missed due to transcriptional inactivation, a genome wide CRISPR activation screen is more desirable. Also, the performance of CRISPR screens is heavily reliant on the quality of genome annotations; continued refinement and upgrade of livestock genome annotations will be crucial for improving efficacy. The current RefSeq annotation for human genome GRCh38.p13 records 5.7 fully supported CDSs per protein coding gene on average whereas the average number included in the cow assembly ARS-UCD1.2 is merely 3. This lack of information might partially explain why our screen did not recapture some of the previously documented host genes such as MAPK8/9, CTNNB1, and SGK1/2/3^42,43,81^. Another factor could be functional redundancy; genes with redundant equivalents can be difficult to discover using single gene targeting knockout libraries. This is perhaps the primary reason why our screen identified all common subunits shared by GARP and EARP i.e. VPS51, VPS52 and VPS53 but not the complex specific subunits VPS50 and VPS54, as our data show that GARP and EARP serve as alternative routes for the virus. A new library design that carefully incorporates gene family and functional redundancy information should further improve the efficacy of genome wide screens.

## Materials and Methods

For more details, please refer to the **Supplementary Materials and Methods**. Briefly, the btCRISPRko.v1 library design was based on RefSeq annotations of assemblies UMD3.1.1 and btau5.01. CRISPRs were chosen to target the first half of common sequences shared by transcription isoforms from 21,165 mostly protein coding genes, with optimal on-target efficiency and specificity estimated by Azimuth 2.0 and CFD^39^. The library was synthesized as 80-mer single strand oligos in the following format: GCAGATGGCTCTTTGTCCTAGACATCGAAGACAACACCG-N20-GTTTTACAGTCTTCTCGTCGC, with N20 standing for 96,000 variable 20bp CRISPR sequences. The oligos were then PCR converted to dsDNA and cloned into pKLV2-U6gRNA5(BbsI)-PGKpuro2ABFP-W^46^ by BbsI digestion and T4 ligation to make the K2g5 library. The other three libraries were cloned using the same protocol but with different vector backbones. The library was packaged into lentivirus in HEK293FT cells by Calcium Phosphate transfection and transduced into ~100 million Cas9+/+; TRIM5 −/− cells at a MOI of 0.3-0.5 with three repeats. Transduced cells were selected with 1.8 ug/ml Puromycin for 7-10 days, while maintaining a coverage of 300-500x. For both screens, 70-100 million passage 4-5 library transduced cells were infected with GFP tagged BHV-1 virus at a MOI of 2. At 10hpi (1^st^ screen) or 8phi (2^nd^ screen), cells were harvested for FACS sort, to obtain fractions of live cells with varied GFP intensity, i.e. GFP negative, GFP low, GFP medium, and GFP High. The viral infection was repeated three times and genomic DNA was extracted from all fractions and all repeats. The CRISPR containing lentiviral sequences were amplified and barcoded by a 2-step PCR with limited cycles and sequenced on a NextSeq 500 machine using a SR75bp High Output kit. Reads for each sample were then trimmed using cutadapt v1.16^82^ and counted based on the CRISPR sequences they contain using MAGeCK v0.5.8^35^. Pairwise comparisons of CRISPR copy numbers between fractions were also completed by MAGeCK, to identify genes with significantly enriched or depleted guides in different fractions post BHV-1 infection. These genes were identified as candidates and those with interesting biology by pathway analysis and literature search were studied further.

## Data and materials availability

Most data generated in this study have been included in the main files or supplementary files. Any requests for additional data or materials such as plasmids, cell lines or cell clones please direct them to.

## Author contributions

W.S.T, S.G.L, C.B.A.W, A.L and R.G.D conceived the study and obtained funding, W.S.T designed and produced the libraries, performed the screens and analyzed the screen data, W.S.T, E.R, and I.D conducted validation experiments, W.S.T, E.R, I.D and R.G.D analyzed the validation data, A.L contributed computing resource, W.S.T wrote the manuscript; All authors reviewed and/or edited the manuscript.

## Acknowledgements

This study is funded by BBSRC grant BB/P003966/1, and BBSRC strategic funding to the Roslin Institute via grants BB/P013740/1 and BB/P013759/1. We thank Dr. Peter Wild and other colleagues at ZTH for sharing the GFP tagged BHV-1 strain, Dr. Kosuke Yusa from the Wellcome Sanger Institute, Amy Tong from Professor Jason Moffat’s lab at the University of Toronto, and Professor Keisuke Kaji from the Scottish Center for Regenerative Medicine at the University of Edinburgh for sharing their protocols, and other members of the CRISPR screening community for wonderful insights and techniques. We are also grateful to the imaging facility staff Dr. Bob Fleming and Graeme Robertson and the CSU unit for support, the virology group for meaningful discussions, and Dr. Lel Eory and Dr. Mazdak Salavati for tips in data plotting, all based at the Roslin Institute. We would also like to extend our appreciation to members of the NGS team, Angie Fawkes, Richard Clark and Lee Murphy at the Edinburgh Clinical Research Facility for sequencing our samples.

