## Supplementary Data for "Genome-wide CRISPR knockout screen reveals membrane tethering complexes EARP and GARP important for Bovine Herpes Virus Type 1 replication"

**References**

**Lentivirus transduction efficiency in wt MDBK cells is very low**

Compared to HEK293FT cells, MDBKs are 30-60 times less transducible (**Fig. S1C, S1D**). Although this is sufficient for single gene knockouts with our efficient serum free lentivirus packaging protocol (**Fig. S1A, 1B**), it becomes labour intensive and costly to deliver a genome wide CRISPR library, as it requires large quantities and high titres of lentivirus stocks to achieve sufficient coverage. It has been reported that TRIM5a inhibits HIV-1 based lentivirus transduction in cows^1,2^. To overcome this low transduction efficiency obstacle, we decided to knock this gene out in the MDBK cells. The bovine TRIM5a gene has been duplicated multiple times during evolution^3^, and we were able to design a pair of high functioning TALENs (pair 1L+1R, **Fig. 2**) specific to TRIM5-3, the third gene in the TRIM5-related gene cluster that is responsible for the anti-retrovirus activity^1,2^. The TALENs cut immediately upstream of the B30.2 HIV-1 capsid recognition domain, destroying TRIM5a activity.

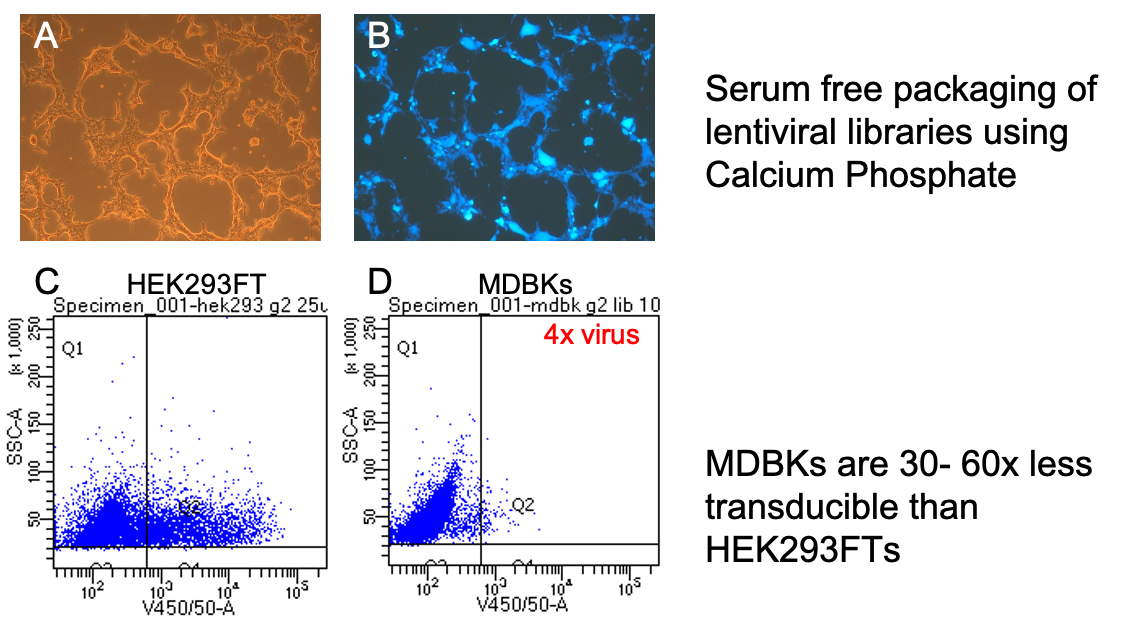

**Supplementary Figure S1. Lentivirus transduction efficiency is low in wt MDBK cells.** A. bright field visualization of HEK293FT cells 48 hours after being transfected with the library plasmid pool, pMD.2 and psPAX2 for serum-free lenti-virus packing. B. Same cells under the BFP filter showing strong BFP expression, indicating efficient transfection and packaging. C. CRISPR library transduction in HEK293FT cells. D. CRISPR library transduction in Cas9+/+ cells with four times the virus.

**Cas9 expression does not affect BHV-1 infection in MDBKs**

To decide whether Cas9 expression would affect BHV-1 replication, we conducted plaque assays in Cas9+/+ MDBKs and wt MDBK cells by infection with the GFP tagged BHV-1 virus (**Fig. S2**). We compared the numbers (**Fig. S2B**) and sizes (**Fig. S2A**) of plaques formed in the three homozygous Cas9 clones, #C, #P and #AH and those formed in the wt MDBK cells. No significant difference between the wt MDBKs and Cas9 clones was observed (p>0.05), indicating that Cas9 expression has no effect on BHV-1 replication in MDBK cells.

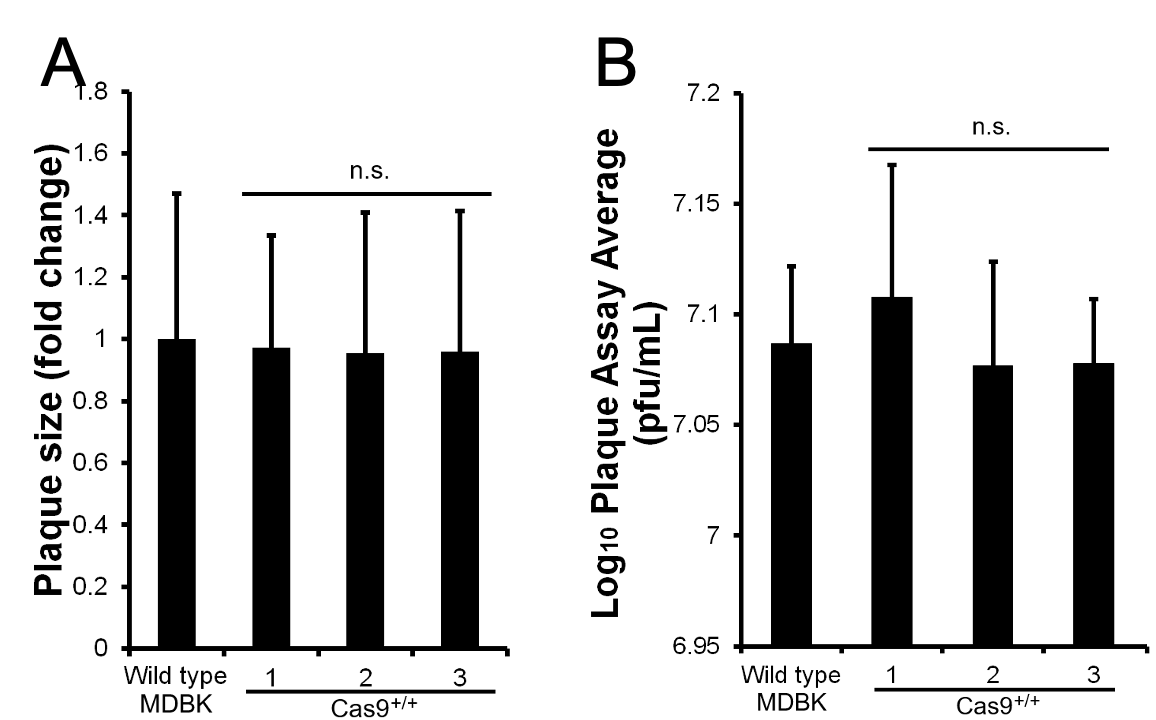

**Supplementary Figure S2. Plaque assays to compare replication of BHV-1 in wild type MDBK cells and homozygous Cas9 clones.** A. Plaque sizes measurements from wt, and three Cas9^+/+^ clones infected with GFP tagged BHV-1 virus, repeat n=3. B. Plaque number counts calculated as pfu/mL from wt, and the Cas9^+/+^ clones.

**The loss of TRIM5a does not affect BHV-1 infection in MDBKs**

To compare BHV-1 replication in wild type MDBKs and TRIM5a knockout clones, we conducted plaque assays using the GFP tagged BHV-1 virus (**Figs. S3, S11**). We studied the numbers (**Fig. S3B**) and sizes (**Fig. S3A**) of plaques formed in wild type MDBKs and two TRIM5-/- clones, A44 and B13. No significant difference between the wt MDBKs and KO clones was evident by pairwise comparisons (p>0.05). We also compared the Cas9+/+ clones to Cas9+/+; TRIM5 -/- clones in terms of plaque sizes and numbers and no difference was observed either (**Fig. S4**), indicating that the loss of TRIM5a has no effect on BHV-1 replication in MDBK cells.

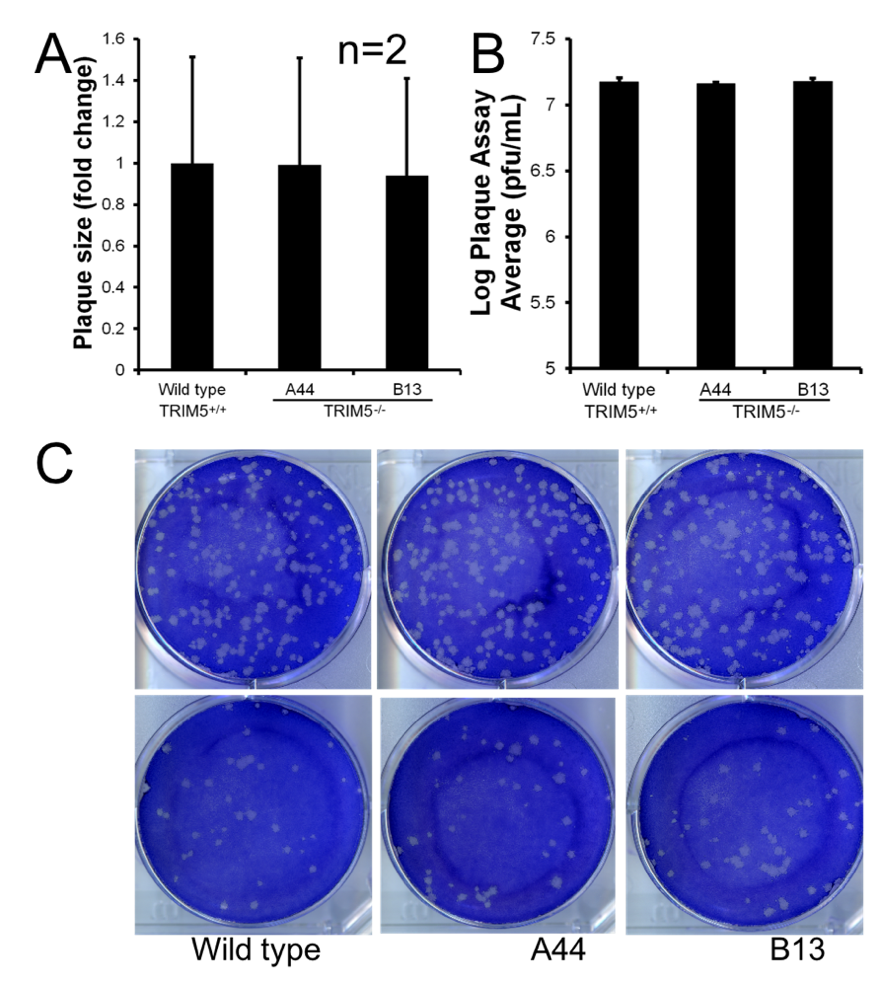

**Supplementary Figure S3. Plaque assays to compare replication of BHV-1 in wild type MDBK cells and TRIM5-/- clones.** A. Plaque sizes measurements from wt, and two TRIM5 KO clones, A44 and B13 infected with GFP tagged BHV-1 virus, repeat n=2. B. Plaque number counts calculated as pfu/mL from wt, and the TRIM5 KO clones. C. Sample plaque assay results with the same quantities of virus.

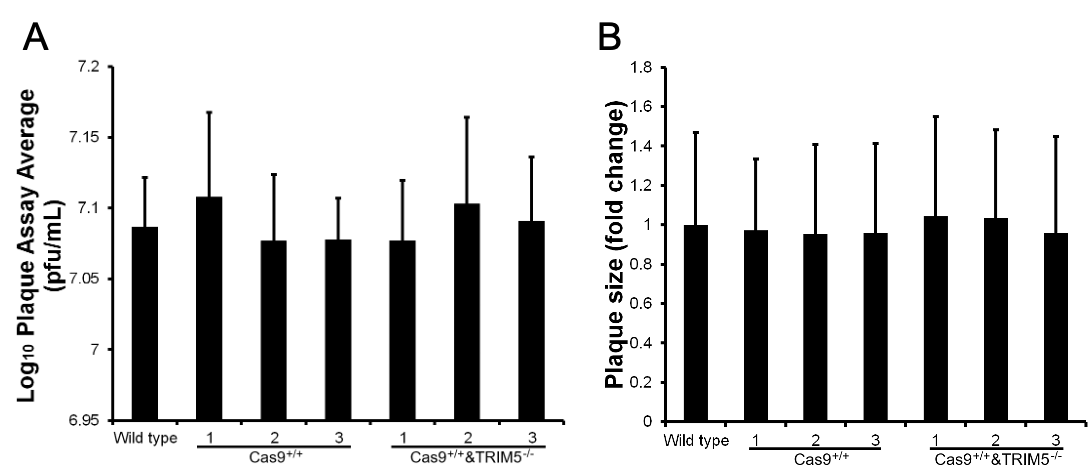

**Supplementary Figure S4. Plaque assay to compare BHV-1 replication in Cas9+/+ only cells and Cas9+/+; TRIM5-/- clones.** A. Titer of virus grown in Cas9+/+ or Cas9+/+;TRIM5-/- cells; B. Average size of plaques grown.

**Library design validation**

Prior to library cloning, its design was validated by testing editing efficiency of guides against a list of genes. Guides were picked randomly from the library for chosen genes and cloned into lentiGuide-Puro or PB_U6gRNA2-CAGpuro for lentiviral or PiggyBac based delivery into Cas9+/+ MDBKs. Alternatively, some guides were *in vitro* transcribed into sgRNA and transfected into MDBKs for leave-no-trace editing. After four days of Puromycin selection or 2-3 days of recovery after transfection with sgRNA, the genomic DNA was harvested for T7 assays and TIDE analysis to determine CRISPR cutting efficiency. Reassuringly, all guides tested are functional with editing efficiency ranging from 11.4% to 79% (**Fig. S5**). And regardless of method, PB and lentivirus delivered similar cutting efficiency (**Fig. S5A**). *In vitro* transcribed sgRNA also mediated efficient editing with up to 65% gene editing (**Fig. S5B**); this selection-marker-free and leave-no-trace editing method is very convenient for generating gene knockout clones.

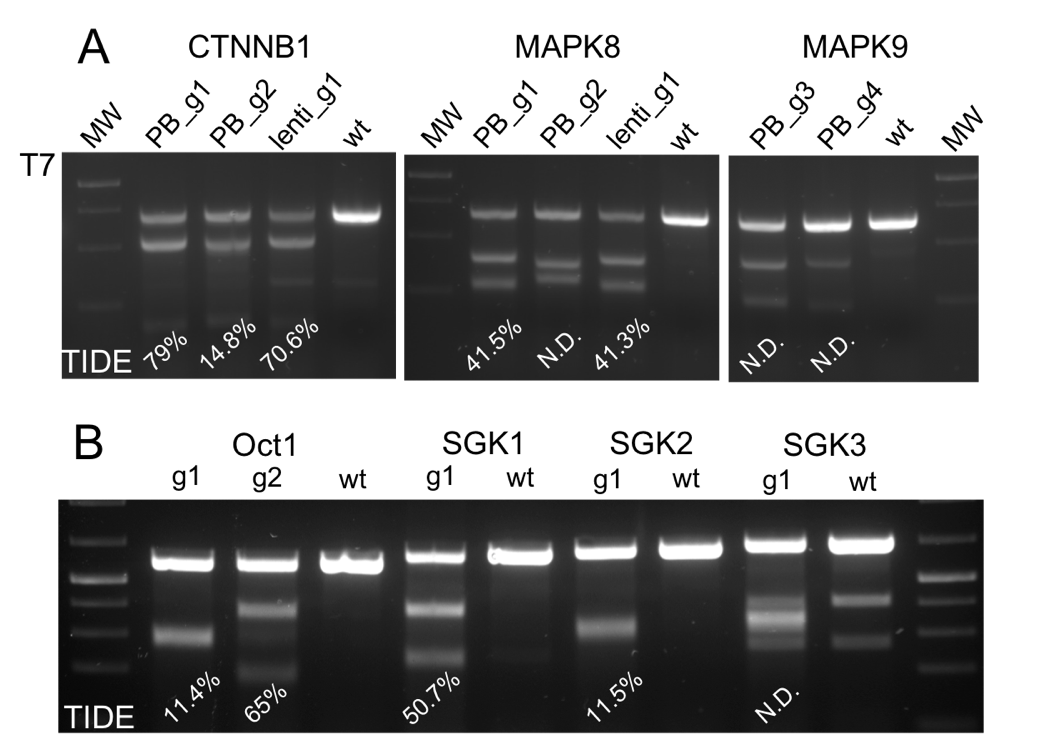

**Supplementary Figure S5. Testing CRISPRs designed for the CRISPRko library in MDBK cells.** A. Cutting efficiency of guides delivered by lentivirus or PiggyBac targeting CTNNB1, MAPK8 and MAPK9. B. Cutting efficiency of guides *in vitro* transcribed as sgRNA and delivered by transfection, targeting Oct1, SGK1, SGK2, and SGK3.

**Cloning, packaging, and transducing lentiviral libraries into Cas9+/+;TRIM5-/- MDBKs**

We produced four genome wide CRISPR knockout libraries for cattle using the same oligo pool (**Table S2**), K2g2, K2g5 (a.k.a btCRISPRko.v1, **Data file 1**), PBg2 and PBg5, based on different sgRNA scaffolds (g2 and g5) and delivery methods (lentivirus and piggyBac). Recently Chen *et al.* ^4^ developed an optimized sgRNA scaffold that was shown to mediate higher CRISPR cutting activity, by removing the potential premature T7 polymerase stop signal and extending the hairpin stem loops for better stability on the target. This has been confirmed in human and mouse cell lines in a few small- and large-scale studies^5,6^. To determine which scaffold we should adopt for our screen, we generated two lentivirus libraries based on these scaffolds (**Fig. S9,10**) and compared their performances in the Cas9 +/+; TRIM5-/- cells (**Fig. 2**). In addition, to make the btCRISPRko.v1 broadly applicable, the CRISPRs were also cloned into PB-U6gRNA(BbsI)-PGKpuro2ABFP and PB-U6gRNA5(BbsI)-PGKpuro2ABFP-W, to generate two additional libraries, PBg2btCRISPRko.v1 and PBg5btCRISPRko.v1 for delivery by transfection and transposition. This approach could facilitate the use of genome wide CRISPR knock out screens in hard to transduce cell types such as primary macrophages.

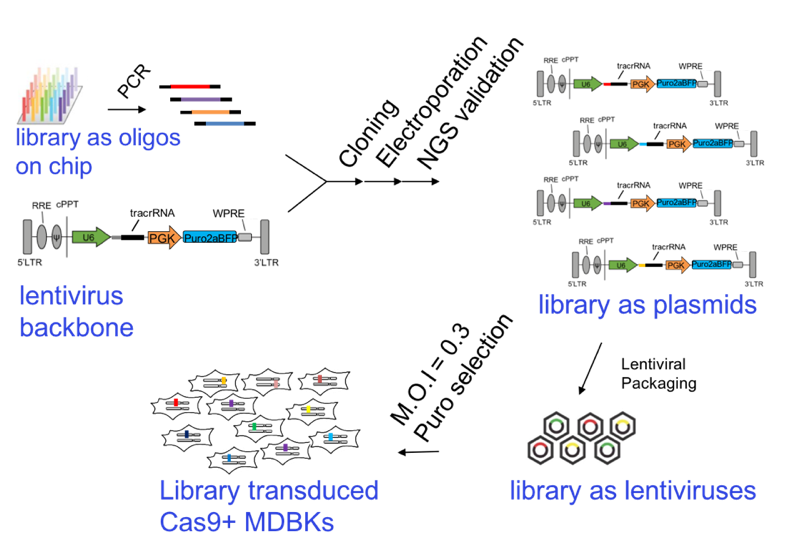

**Supplementary Figure S6. Stepwise CRISPR library cloning, packaging, and transduction.** Library was synthesized as a pool of oligos and PCR amplified to convert to dsDNA. The fragments containing CRISPRs are then ligated into either lentivirus (illustrated as example) or piggyBac vectors that contain a hU6 promoter to drive sgRNA expression and a Puro2aBFP marker for Puromycin selection. If using lentiviral delivery, the library is packaged as lentivirus and transduced into Cas9 expressing cells at low MOI to produce library expressing cells with single sgRNA integrations.

**PiggyBac CRISPR Knockout libraries**

The cloning of these PiggyBac libraries underwent the same quality control procedures as the lentiviral libraries, with above 1,000x depth, below 0.3% background and above 90% accuracy. A colony formation assay was conducted to test whether the libraries could transpose efficiently. When a plasmid expressing the piggyBac transposase i.e. hypBase^7^ was added in a pilot library transfection and selection experiment using Cas9 expressing MDBKs (**Fig. 2**), we obtained significantly more clones after puromycin selection compared to cells without transposase, indicating efficient CRISPR library integration by transposition.

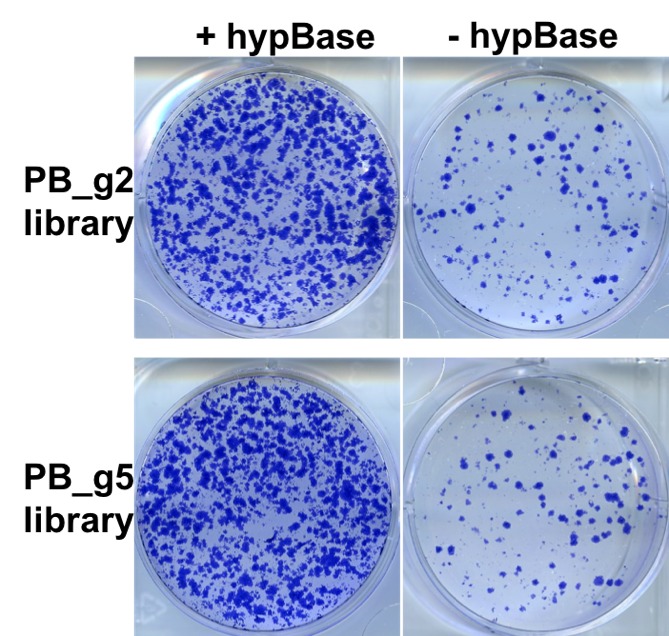

**Supplementary Figure S7. PiggyBac transposition mediated CRISPR library delivery into Cas9+/+ MDBK cells.** Cas9+/+ transfected with either library with (left panels) or without (right panels) transposase hypBase and selected with Puromycin to test library transposability. After Puro selection, cells were stained with Giemsa for colony visualization.

**NGS and pairwise comparisons to assess library performance and identify candidate host genes**

By PCR and next generation sequencing, we obtain copy numbers of guides in all the samples collected from our screen (**Data file 2,3,S1**). Pairwise comparisons between samples can identify enriched or depleted guides, and depletion or enrichment of corresponding genes. To illustration, guide RNA copy numbers are compared between the GFP Negative and GFP High sub-populations, enrichment of guides in the Negative sample relative to the High leads to depletion of genes targeted by these guides in the Negative sample, and identification of essential or pro-viral host genes for BHV-1 infection. On the contrary, depletion of guides in the Negative sample leads to enrichment of genes that can be anti-viral.

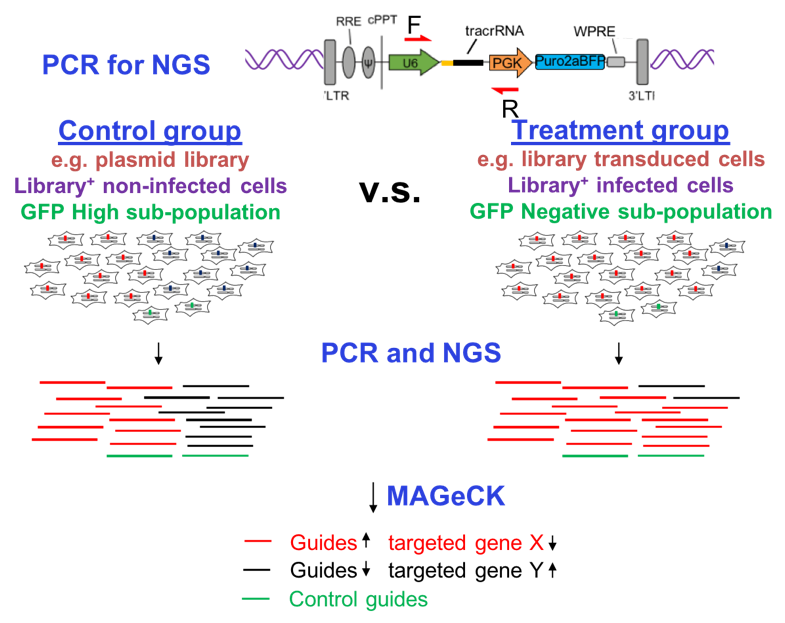

**Supplementary Figure S8. Next generation sequencing of screening samples to identify candidate genes with depleted or enriched guide RNA.** To determine copy numbers of guides, PCR using primers PCR1_Fx + PCR1_R (red arrows, Supplementary Table S5) are used to amplify the fragments containing CRISPR sequences. PCR products from different samples are then barcoded using a 2^nd^ PCR with a unique combination of indexes and sequenced as a pool. Comparison of guide RNA copy numbers between samples is conducted by MAGeCK to identify enriched or depleted guides and their targeted genes.

**Quality control and performance comparison between the lentiviral g2 and g5 libraries**

To examine guide RNA distribution and accuracy, the two lentiviral libraries were sequenced by NextSeq at 40x sequencing depth (**Data file S1**). After read processing and counting all guides included in the libraries, we observed good distribution and presence of the majority of the guides for both libraries (99.6% vs 98.8%). However, the g5 library is slightly less uniform than g2, likely due to PCR artefacts during library cloning. After deciding on using the lentiviral g5 library for our screening, we re-sequenced the K2g5 library with higher sequencing depth, at 200x together with library transduced cells prior to our BHV-1 screen (**Data file 2**).

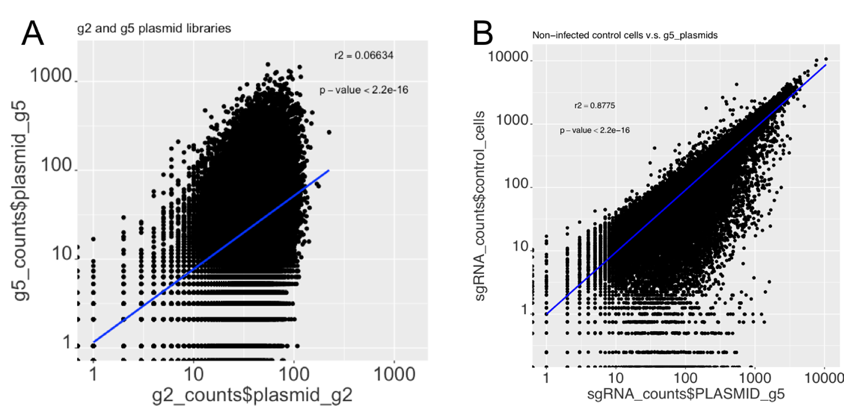

**Supplementary Figure S9. NextSeq of the CRISPR libraries to examine CRISPR copy number distribution.** A. Sequencing of the K2g2 and K2g5 plasmid libraries at 40X sequencing depth. B. Sequencing of the K2g5 library and library transduced Cas9+/+; TRIM5-/- cells at 200X sequencing depth.

By converging data from several CRISPR screening studies, Hart *et al.* identified a list of 684 core essential genes (CEG2.0) shared among 17 genome wide CRISPR knockout screens in human cell lines^8^. The CEGs can be used as good indicators of library performance; since if a CEG targeting CRISPR works well, it should be gradually removed from the cell population during extended culture. Thus, a collective drop out of CEG2.0 targeting guides indicates library functionality; and the bigger the shift, the better the performance. To compare performance of the two lentivirus libraries, we packaged, titrated, and transduced them into MDBK cells using the same protocols. After seven days of Puromycin selection, genomic DNA were harvested from the two cell populations and the CRISPR regions were PCR amplified and sequenced by NextSeq at 40x depth (**Fig. S8, Data file S1**). We examined the degrees of drop out of CEG2.0 targeting guides in the two MDBK populations relative to the plasmid libraries and saw a much bigger shift in g5 transduced cells compared to g2, indicating better performance of g5 than g2 (**Fig. S10**). For both libraries, we observed relative stable or unchanged distribution of guides targeting non-essential genes and non-cutting control guides, indicating good overall specificity and control of off targeting from both libraries. Based on these results, we decided to use g5 library transduced cells for our BHV-1 screens.

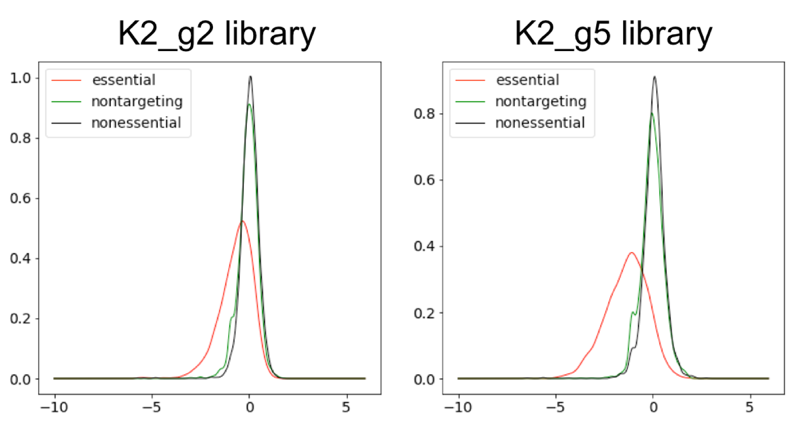

**Note: re-plot to have the same y-axis scale for better comparison!!! Also, calculate percentages!**

**Supplementary Figure S10. Library performance and specificity comparison between the K2g2 and K2g5 libraries.** Plots show log2 fold changes in copy numbers of guides targeting essential genes(red), non-essential genes(black), and non-targeting control guides(green) in the two cell populations transduced with the K2g2 or K2g5 library compared to the plasmids.

**GFP tagged BHV-1 virus replicates in MDBKs**

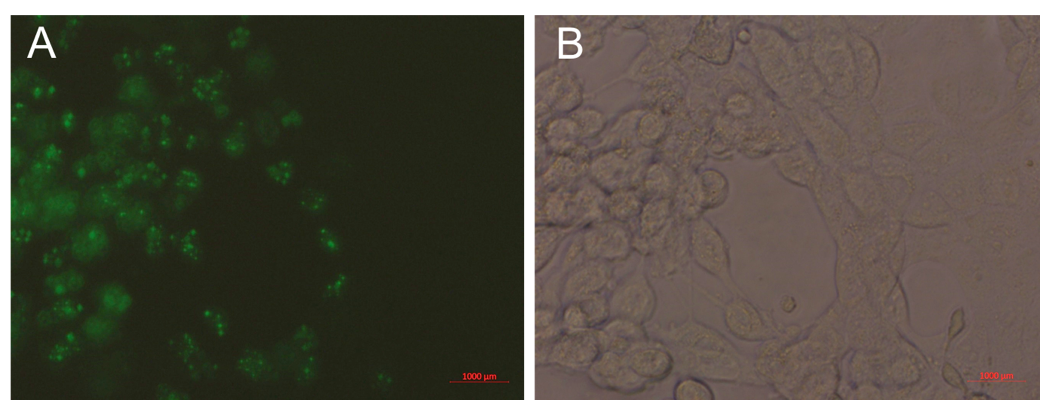

**Supplementary Figure S11. GFP tagged BHV-1 virus infecting wt MDBK cells.** A. 8 hours post infection of MDBKs by the GFP tagged virus. B. image of the same cells under bright field.

**FACS sort to isolate sub-populations with varied intensities of GFP**

For the 1^st^ screen, 9.5-10 hours post infection of library transduced TRIM5-/-; Cas9+/+ MDBKs at MOI=2, cells were harvested for FACS sorting to isolate sub-populations with different levels of BHV-1 infection based on GFP intensity, Negative, Low, Medium, and High (**Fig. S12**). The gating for the Negative was based on the non-infected cells, and the Low, Medium and High gates have equal width across the spread of the rest of the cells. Genomic DNA is then isolated from these sub-populations, PCR amplified and sequencing by Illumina NextSeq to examine distribution of guides (**Fig. S8**, **Data file 2**). For the 2^nd^ screen, cells were infected the same as the 1^st^ screen but harvested at 8 h.p.i. The gatings were set to collect all GFP Negative cells based on the non-infected cells and ~10% of live cells per sub-population for GFP Low, Medium and High with substantial space between gates. Cells were then processed and sequenced as the 1^st^ screen.

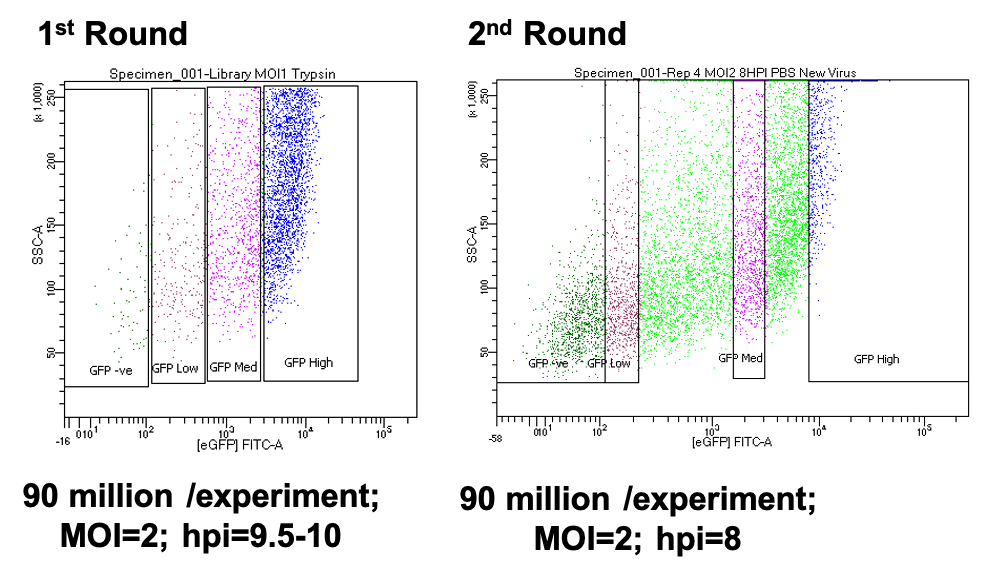

**Supplementary Figure S12. FACS sort to collect sub-populations of different degrees of BHV-1 replication.** FACS plots with four gatings to collect sub-populations of live cells with GFP Negative, GFP Low, GFP Medium, and GFP High signals from 1^st^ or 2^nd^ round of screen.

**CRISPR screen identifies pro-viral genes**

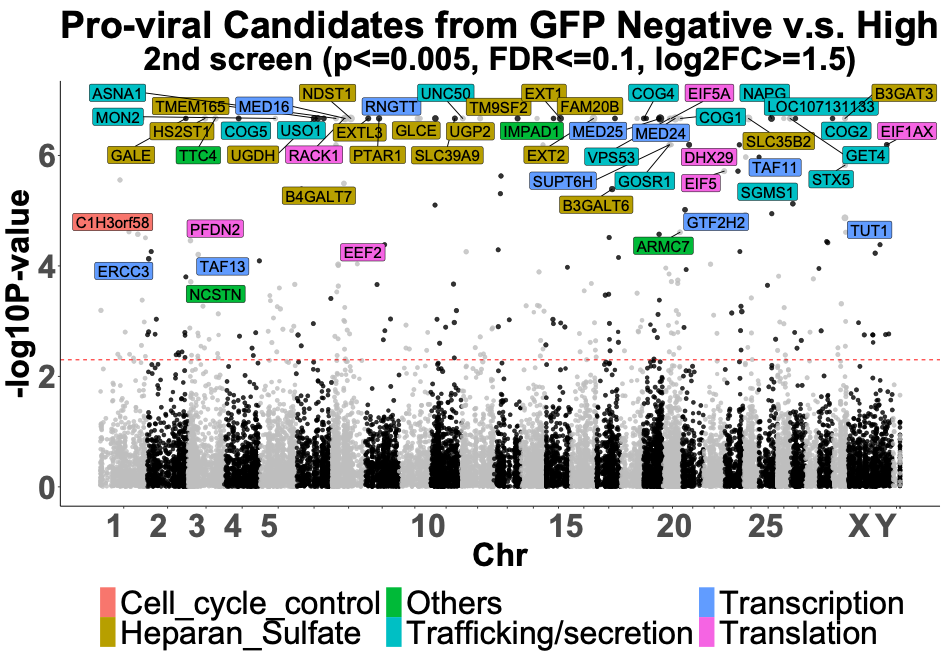

**Supplementary Figure S13: Candidates identified in the GFP Negative sub-population.** Data shown is from 2^nd^ screen, labels only applying to pro-viral candidates with stringent cut-offs: p<0.005, FDR<=0.1, and log2fc>=1.5 (**Data file 3**).

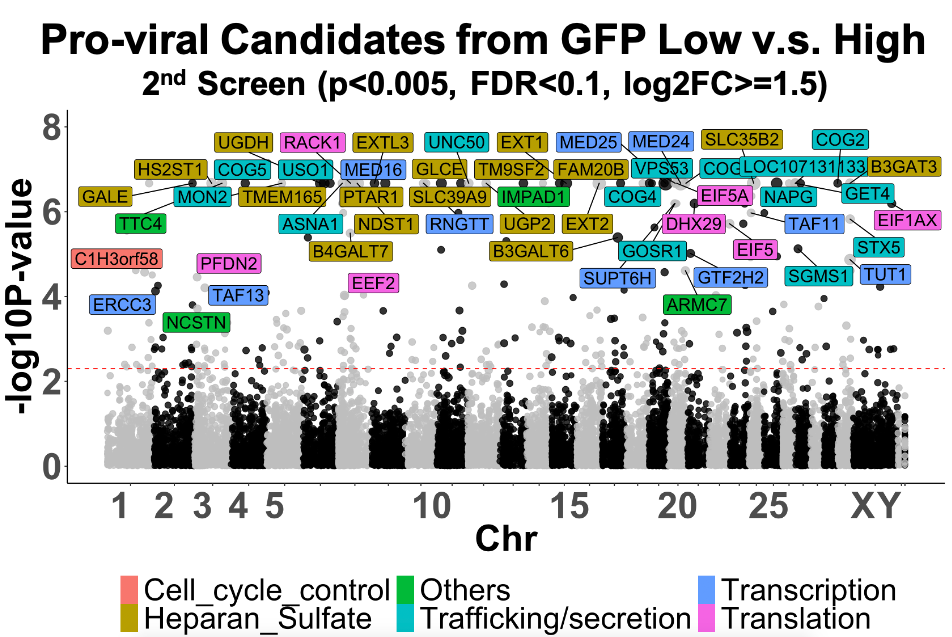

**Supplementary Figure S14: Candidates identified in the GFP Low sub-population.** Data shown is from 2^nd^ screen, labels only applying to pro-viral candidates with stringent cut-offs: p<0.005, FDR<=0.1, and log2fc>=1.5 (**Data file 3**).

**The loss of GARP and EARP also severely affects AIHV-1 replication in MDBK cells**

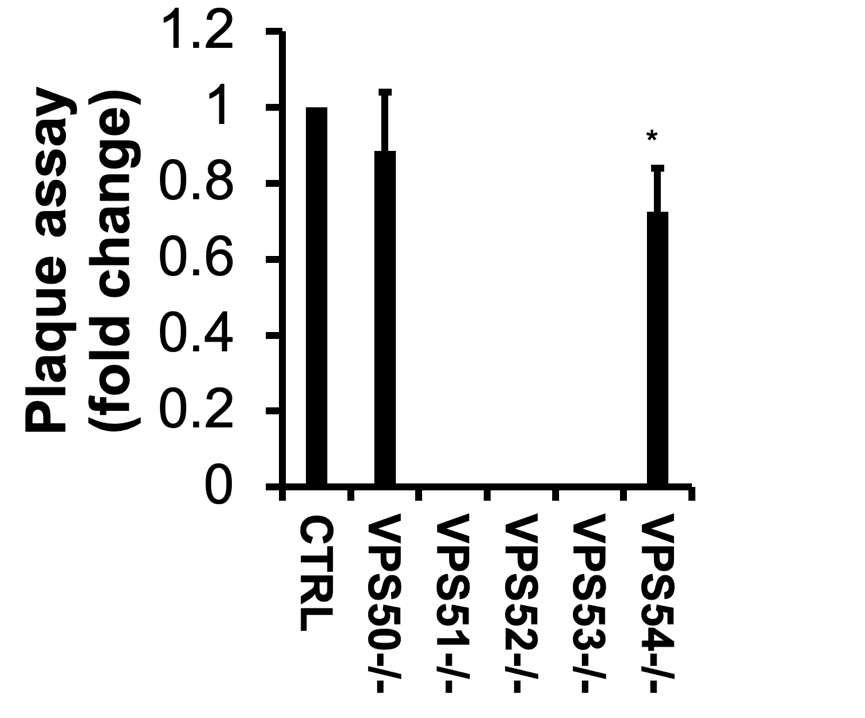

**Supplementary Figure S15: AIHV-1 replication in VPS52 KO.** Results of plaque assays conducted on Cas9+/+ MDBKs (CTRL) or MDBK clones with VPS50, VPS51, VPS52,VPS53, and VPS54 bi-allelic KO. Titers shown are relative to Cas9+/+ with result from Cas9+/+ set as 1. n=3, *: p<0.05 between CTRL and VPS54-/-.

**The loss of GARP and EARP reduces viral genomic DNA in MDBKs after BHV-1 infection**

A

B

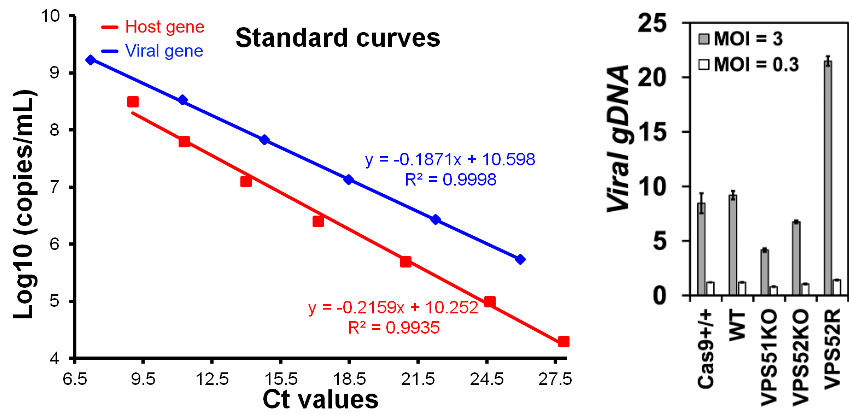

**Supplementary Figure S16. qPCR quantification of viral genomic DNA in MDBK cells infected with BHV-1 at 6 h.p.i. A.** Standard curves used to calculate copy numbers of host genome or viral genome based on CT values. Host gene PVRL2 was used as internal control; PCR primers against viral gene ICP4 was used for viral gDNA quantification. **B.** relative copy numbers of viral genomes in Cas9+/+ and WT control cells, VPS51KO and VPS52KO cells and VPS52R rescue cells. Values are expressed relative to that of Cas9+/+ which was set at 1. Note: data show here was obtained from a single experiment with three technical repeats.

**Supplementary Table S1. Design statistics of the btCRISPRko.v1 library**

| **Total genes targeted** | 21,165 | **No. genes with 3 guides** | 112 |
| --- | --- | --- | --- |
| **Total targeting guides** | 94,000 | **No. genes with 2 guides** | 71 |
| **No. genes with 6 guides** | 2 | **No. genes with 1 guide** | 38 |
| **No. genes with 5 guides** | 9,704 | **Control guides** | 2,000 |
| **No. genes with 4 guides** | 11,238 | **Total guides** | 96,000 |

**Supplementary Table S2. Libraries produced from this study**

| **library** | | **Delivery**  **method** | **scaffold** | **colony-based QC** | | **Sanger QC**  **% correct** | **NextSeq-based QC** | |
| --- | --- | --- | --- | --- | --- | --- | --- | --- |
|  |  |  |  | **depth** | **background** |  | **coverage** | **% correct** |
| K2g2 | lentivirus | | Zhang^9^ | 1361x | 0.6% | 21/22(95.5%) | > 99.70% | N.D. |
| K2g5 | lentivirus | | Chen^4^ | 1300x | 1.5% | 22/23(95.7%) | > 99.9% | 96.02% |
| PBg2 | piggyBac | | Zhang | 4700x | 0.2% | 24/24(100%) | N.D. | N.D. |
| PBg5 | piggyBac | | Chen | 1270x | 0.3% | 21/23(91.3%) | N.D. | N.D. |

**Notes:** K2g2: lentivirus library with the original sgRNA scaffold; K2g5: lentivirus library with the optimized sgRNA scaffold; PBg2: PiggyBac library with the original sgRNA scaffold; PBg5: PiggyBac library with the optimized scaffold; N.D.: not determined.

**Supplementary Table S3. Number of cells recovered from 1^st^ screen**

|  | **GFP Negative** | **GFP Low** | **GFP Medium** | **GFP High** | **Non-infected** |
| --- | --- | --- | --- | --- | --- |
| **Repeat 1** | **1.07** | **3.88** | **5.47** | **11** | **50** |
| **Repeat 2** | **1.2** | **3.84** | **6** | **15.7** | **50** |
| **Repeat 3** | **0.53** | **1.11** | **2.59** | **6.76** | **50** |
| **Repeat 4** | **1.07** | **3.6** | **7.01** | **23** | **50** |

**Note: Cell numbers are in Millions.**

**Supplementary Table S4. Number of cells recovered from 2^nd^ screen**

|  | **GFP Negative** | **GFP Low** | **GFP Medium** | **GFP High** | **Non-infected** |
| --- | --- | --- | --- | --- | --- |
| **Repeat 1** | **1.66** | **3.1** | **2.53** | **2.45** | **70** |
| **Repeat 2** | **4.3** | **4.3** | **3.5** | **3.3** | **70** |
| **Repeat 3** | **3.6** | **5.4** | **4.9** | **3.7** | **70** |
| **Repeat 4** | **2.14** | **3.0** | **3.27** | **4.66** | **70** |

**Note: Cell numbers are in Millions.**

**Supplementary Table S5. Knockout clone genotypes**

| **Purpose** | **Clone I.D.** | **Genotype** |
| --- | --- | --- |
| TRIM5a^-/-^; Cas9^+/+^ | 1B2 | -13/-13 |
| TRIM5a^-/-^; Cas9^+/+^ | 1C1 | +2/+2 |
| TRIM5a^-/-^; Cas9^+/+^ | 2A5 | -2/-10 |
| TRIM5-/- | A44 | -1/-1 |
| TRIM5-/- | B13 | -29/-35 |
| VPS52KO | B1 | -8/-11 |
| VPS52KO | B4 | -5/-10 |
| VPS52KO | C1 | +8/+8 |
| VPS52KO | C4 | -8/-8 |
| VPS51KO | A9 | -1/-2 |
| VPS51KO | B7 | -5/-5 |
| VPS51KO | C2 | -2/+1 |
| VPS51KO | C3 | +1/+1 |
| VPS53KO | E2 | -5/-7 |
| VPS53KO | E5 | -5/-7 |
| VPS53KO | F11 | -2/-2 |
| VPS53KO | F3 | -2/-17 |
| VPS54KO | A4 | +1/-7 |
| VPS54KO | C6 | -5/-5 |
| VPS54KO | D5 | -5/-10 |
| VPS54KO | D7 | +1/+1 |
| VPS50KO | C3 | -1/+25 |
| VPS50KO | A4 | -950/-950 |
| VPS50KO | D4 | -950/-950 |
| VPS50KO | E4 | -950/-950 |
| VPS50;54dKO | C3A2 | -1/+25;-2/-2 |
| VPS50;54dKO | D4B4 | -950/-950;-7/-7 |
| VPS50;54dKO | D4F3 | -950/-950;-7/-8 |
