## Supplementary Materials and Methods for "Genome-wide CRISPR knockout screen reveals membrane tethering complexes EARP and GARP important for Bovine Herpes Virus Type 1 replication"

**References**

**Genome-wide library design**

**Step 1**: **Extract common coding sequences among CDS isoforms**

The latest RefSeq annotations for the bovine genome assemblies (as of Nov. 2017) GCF_000003055.6 from UMD3.1.1 and GCF_000003205.7 from btau5.01 were downloaded from the FTP of NCBI, and the Y chromosome from btau5.01 was combined with UMD3.1.1. For each protein coding gene, a python script utilizing the “intersect” sub-command from the bedtools v2.26.0 suite was used to extract coordinates of shared coding sequences among CDS isoforms.

**Step 2: Extract candidate CRISPRs**

These coordinates were then used to extract genomic sequences from the assembly using the bedtools v2.26.0 “getfasta” sub-command. All 20bp sequences immediately upstream of “NGG” tri-nucleotides on both the “+” and “-“ strands were extracted but only guides that met the following criteria were included in the candidate list: **a**. have a 20-80% GC content; **b**. do not contain BbsI binding sites “GAAGAC” and “GTCTTC”, and **c**. do not contain any of the following sequences “N”, “AAAA”, “TTTT”, “GGGG”, and “CCCC”. For each candidate, the cutting position relative to the common coding sequence was used to calculate percentage peptide.

**Step 3: Estimate on-target cutting efficiency**

For every candidate along with its percentage peptide, the on-target efficiency was estimated on the Eddie supercomputing cluster using Azimuth 2.0 developed by Microsoft and the Broad Institute^1^. For each gene, all guides were then ranked based on their cutting efficiency from high to low.

**Step 4: Extract off-target sites and estimating off-targeting efficiency**

On the Eddie cluster, the fasta sequences for UMD3.1.1 and the Y chromosome from btau5.01 were indexed by the bwa 0.7.12-r1039 “index” sub-command. All CRISPR candidates were then aligned back to the indexed genome using the “aln” sub-command to identify and extract 20bp sequences with up to three mismatches to the candidate. Only sequences immediately upstream of “NGG”, “NAG” and “NCG” were regarded as potential off-targets and the cutting efficiency at every off-target relative to the on-target was estimated by the CFD package, also developed by Microsoft and the Broad Institute^1^.

**Step 5**: **Select guides**

For each gene, between 4 to 5 guides were selected for inclusion in the final library by processing the ranked list from Step 3 following these rules: **1**. they cut closest to the 5’; **2.** They had the highest on-target efficiency; and **3.** they had the least off-targets with CFD > 0.2 that reside in exons. For some genes, multiple rounds of selection with gradually relaxed rules were needed to identify sufficient numbers of guides; the final list of targeting guides contained 94,000 guides in total.

**Step 6: Generate non-cutting control guides**

A custom python script was used to generate one million random 20bp sequences but only those that met all criteria specified in Step 2 were selected as potential control guides. Sequences selected were also aligned back to the genome as described in Step 4 and only those that align to one or two non-exon sequences with CFD scores less than 0.05 were included in the final 2,000 control-guide list.

**Step 7**: **Synthesize the library as single strand oligos**

The targeting and non-targeting guide sequences were combined, and adapter sequences added to the 5’ and 3’ ends as follows: 5’- GCAGATGGCTCTTTGTCCTAGACATCGAAGACAACACCG-N_20_-GTTTTAGTCTTCTCGTCGC -3’. The list of 96,000 oligos was sent to Twist Bioscience (San Francisco, California) for synthesis and the oligos were delivered as a desalted pool in a single tube.

**Library cloning**

**Genome-wide library cloning:** The pool of 80-mer oligos was PCR amplified to produce full length dsDNA segments using primer set 79-mer-L1+79-mer-U1 for cloning the g2 library and set 79-mer-L1v2+79-mer-U1 for the g5 library ( see Table S5 for primer sequences) . Purified CRISPR containing PCR products were then ligated into a linearized vector, pKLV-U6gRNA (BbsI)-PGKpuro2ABFP (a gift from Kosuke Yusa)^1^ for the K2g2 library or pKLV2- U6gRNA5(BbsI)-PGKpuro2ABFP-W^2^ for the K2g5 library (Addgene ID:67974), using BbsI-HF and T4 ligase. The PBg2 and PBg5 libraries were cloned using the same protocol. These vectors contain a hU6 promoter to drive CRISPR expression, a selection marker Puro_2A_BFP for titration and enrichment of transduced cells, and lentiviral elements for viral packaging or PiggyBac repeats for transposition. Ligated products from 12 parallel reactions were pooled, cleaned and electroporated into electro-competent cells four times. After one hour of recovery at 37° Celsius, 5ul of bacteria were used for plating on Agar plates with 100ug/ml Ampicillin with serial dilutions to estimate coverage. The bulk remaining culture was maintained in 500ml 2XLB with 100ug/ml Ampicillin and plasmid maxipreps were prepared from the overnight culture using the Qiagen Maxi plus kit. The depth, coverage and accuracy of the libraries was determined using the Illumina NextSeq 500 with the SR75 high output kit.

**Tissue culture**

MDBK cells were maintained in DMEM culture media supplemented with 2% or 5% horse serum, 0.5% Pen/Strep, 1% L-Glutamine, 1% Sodium Pyruvate, and 1% NEAA, in a 37°°C incubator with 5% CO_2_. When confluent, cells are passaged 1:6 or 1:9 using Trypsin and they become confluent in 2-3 days. Cells were frozen at concentrations between 1-5x 10^6^ /ml in freezing media made of 10% TC grade DMSO and 90% horse serum or culture media and transferred to -150° Celsius after overnight storage at -80° Celsius. HEK293FT cells are maintained in DMEM supplemented with 10% fetal calf serum and 0.5% Pen/Strep, in a 37°°C incubator with 5% CO_2_. When confluent, cells were trypsinized for 1 minute and expanded 1:9 for subsequent lentivirus packaging.

**Transfection**

0.5-2 x10^6^ MDBK cells were transfected with plasmid DNA, TALEN mRNA, or sgRNA using a Neon electroporator with the following program: 1200v, 30ms, 2 pulses.

For Cas9 targeting to rosa26, 1ug of TALEN mRNA pair #1.6 and 5ug of HDR template were co-transfected into wt MDBKs. After 2 days of incubation at 33°°C, cells were recovered for one day at 37°°C before dilutional cloning and colony isolation.

For TRIM5a knockouts, 1ug of TALEN mRNA pair were transfected into wt MDBKs or Cas9 +/+ MDBKs. Cells were cultured and recovered under the same conditions as the Cas9 targeting experiment prior to colony isolation.

For generating knockout clones using CRISPR sgRNA, 1ug *in vitro* transcribed sgRNA was transfected into Cas9 +/+ MDBKs and cells recovered at 37°°C for two to three days prior to dilutional cloning. For knockouts using PiggyBac transposons, 2.5ug of PiggyBac plasmids carrying hU6 promoter driven gene specific sgRNAs was mixed with 1ug of pCMV-hypBase transposase for co-transfection into Cas9 +/+ MDBKs.

To create rescued cell lines, VPS51, VPS52, and VPS53 KO clones are co-electroporated with a PiggyBac vector carrying the cDNA expression cassettes and pCMV-hypBase supplying PBase at the ratio of x ug : y ug per transfection.

**Serum free lentivirus library packaging, titration and transduction**

Once passing quality control, the lentivirus library was packaged in HEK293FT cells using the Calcium Phosphate transfection method (Supplementary Figure S2A and S2B). One day prior to transfection, 20x 15cm petri dishes were each seeded with 5 x106 early passage HEK293FT cells. 2 hours before transfection on the second day, cells were fed with 10 ml prewarmed fresh media for each plate. Cells were then transfected, using a well-established Calcium Phosphate protocol^2^, with the CRISPR library plasmid pool and the two packaging plasmids pMD.2 and psPAX2. The day after transfection, cells were fed with 20ml serum free media buffered with HEPES. 24 hours after media change, the supernatant with cell debris from each plate was combined, cleared by low speed centrifugation at 300xg for 5 minutes and filtered through a 0.45um low protein binding membrane. The lentivirus stock was then aliquoted into 50ml conical tubes and stored at -80 Celsius until the day of library transduction.

For titration, 3x10^5^ MDBK cells were plated on one 6-well plate 10 hours prior to transduction. 100ul, 80ul, 40ul, 20ul, 10ul or 0ul lentivirus supernatant were mixed separately with 0.8ug/ml Polybrene in culture media with 1ml in total volume in 1.5ml Eppendorf tubes. The culture media was then aspirated from the 6-well plate and the lentivirus dilutions added to the cells immediately. 24 hours after incubation, the virus was then removed and 2ml of fresh media added to each well. The cells were cultured for another day prior to FACS to detect percentage of cells with BFP. The quantity of virus used for the well with ≤30% transduced cells was chosen to calculate the amount of virus needed for the screen at a MOI=0.3.

To produce library transduced cells for the screen, the lentivirus supernatant was used for transduction without prior concentration. 10 hours prior to transduction, a total of approximately 9x 10^7^ TRIM5 -/-; Cas9+/+ MDBKs were plated in 12X T175 flasks. Cells were then transduced at MOI of ~ 0.3 by diluting viral stocks into 10ml total volume using culture media containing 2% horse serum and 0.8ug/ml Polybrene (final concentration). 24 hours after transduction, the virus was replaced by fresh media with 2% horse serum. 48 hours after transduction, the media was replaced with media containing 1.8ug/ml Puromycin for 8-10 days to obtain a pure population of transduced cells. The transduction was repeated three times and cells from every passage were frozen in culture medium supplemented with 10% DMSO and stored at -150° Celsius. To ensure that the libraries are representative and that all the sgRNAs are represented, genomic DNA is isolated from passage 4 transduced cell pools and using primers directed against the lentiviral construct flanking the sgRNA sequences (Supplementary Figure S14), the inserted sequences are amplified and sequenced, providing counts of sgRNA occurrences.

**TALEN assembly and *in vitro* transcription of TALEN mRNAs**

TALENs were assembled the using Golden Gate TALEN and TAL Effector Kit 1.0 following the published protocol^3^. In place of the destination vectors included in the kit, PMC-DeltaTAL was used in the second reaction. This plasmid was constructed by transferring the DeltaTAL fragment in between restriction sites^4^ into the multiple cloning site of pMC128^5^. TALENs were sequence confirmed following bacterial PCR. To transcribe the TALEN DNA pairs into mRNA, 10ug of the left TALEN was combined with 10ug of the right TALEN and linearized overnight with 20U of NotI-HF. The digest was treated with RNAsecure and cleaned up by the MinElute PCR cleanup kit. 1ug eluted linear DNA was then used for *in vitro* transcription and polyA tailing with the HiScribe T7 ARCA mRNA Kit. The polyA mRNA products were purified using a RNeasy column, nanodrop tested for concentration, aliquoted and stored in -80°C.

**Individual sgRNA cloning and *in vitro* transcription of sgRNAs**

Individual guides were cloned into either PB_U6gRNA5-PGKpuro2ABFP-W or pKLV2-U6gRNA5(BbsI)-PGKpuro2ABFP-W by BbsI digestion and T4 ligase ligation as published^6^.

To generate knockout clones by transfecting sgRNA, sgRNA was synthesized by *in vitro* transcription using the HiScribe T7 Quick High Yield RNA synthesis kit. Briefly, to prepare the DNA template for transcription, a PCR was carried out using ttaatacgactcactatagGN19GTTTAAGAGCTATGCTGGAAAC as forward primer, AAAAGCACCGACTCGGTGCC as universal reverse primer and pKLV2-U6gRNA5(BbsI)-PGKpuro2ABFP-W serving as PCR template. The forward primer contains a T7 promoter (lower case), G plus last 19 bp of desired CRISPR sequence (GN19), and partial sequence of the g5 scaffold (underscored sequence). The PCR product was purified using the MinElute PCR cleanup kit and the T7 promoter in the ~110bp amplicon used to transcribe the downstream sgRNA. The *in vitro* transcription was done according the kit manual with 100ng PCR product as template and overnight incubation. The reaction was purified with a RNeasy column, and the sgRNA tested for concentration using a nanodrop spectrometer^7^, aliquoted and stored at -80°C.

**Gene editing efficiency testing by T7 and TIDE**

Two to three days after transfection with TALEN mRNA or individual CRISPR sgRNAs, crude genomic DNA was extracted from cells using QuickExtract DNA Extraction Solution following the product manual. The extract was diluted 1:10 in nuclease free water and 1ul of the dilution used for PCR using primers flanking the target site and Phusion HF polymerase with supplied buffer, producing amplicons between 300-800bp in size. 2ul of PCR reaction is ran on 2% Agarose gel to estimate amplicon concentration. Without purification the calculated volume of PCR reaction containing ~200ng PCR product based on the agarose gel was denatured at 95°°C for 5 minutes and re-annealed by dropping the temperature to 25°C at 0.1°C per second in a thermo cycler, at the end of the program the temperature was reduced to 4°C. Right after re-annealing, 1ul T7 Endonuclease I was added to the PCR reaction directly and incubated at 37°C for 30 minutes. Immediately after incubation, the reaction was analyzed on a 2% Agarose gel. The gel image was processed by the Densitometry function of ImageJ and the percentage editing was calculated as described previously^4^. The editing efficiency was also examined by TIDE analysis (<https://tide.deskgen.com/> )^8^ following Sanger sequencing of purified PCR products.

**Generating the Cas9+/+, and Cas9+/+; TRIM5 -/- clones for the screen and gene editing**

To generate Cas9+/+ clones, TALEN pair TAL1.6 was chosen out of all TALENs and CRISPRs designed that target first intron of the rosa26 locus. It cuts with 52% cutting efficiency based on T7E1 assay and 1ug of the mRNA was co-transfected with 5ug plasmid pMT2.0_bovRosa26_EF1a_Cas9_blast using Neon. After 2 days of incubation at 33°C, cells were recovered at 37°C for two days and then selected with 10mg/ml Blasticidine for four days. Single cell clones were isolated by dilution cloning and genotyped by PCR, homozygotes were expanded and stored in -150°C.

To generate Cas9+/+;TRIM5-/- clones, the population of cells after Blasticidine selection from above was transfected with 1ug of TAL1L+1R mRNA, then cultured and recovered as above. New clones were isolated by dilution cloning and they were screened using a GFP expressing lentivirus. Only those that appeared to have higher % of GFP+ cells judged by eyes were genotyped for both Cas9 KI and TRIM5a KO by PCR and sequencing. Clones homozygous for both Cas9 KI and TRIM5a KO (Cas9+/+;TRIM5-/-) were screened with GFP lentivirus again and the transduction efficiency was measured by FACS to confirm phenotype. Only those with the best morphology and confirmed with enhanced transduction efficiency were expanded for the screen.

**Colony isolation by dilutional cloning and genotyping**

For plasmid-based gene targeting, cells are selected for four days with 10mg/ml Blasticidine or 1.8ug/ml Puromycin depending on the construct. After drug selection, cells are plated at 50-100 cells per dish on 10cm dishes for colony formation. After ~7 days of incubation, colonies are picked, expanded and genotyped by Sanger sequencing and TIDE analysis. For TALEN mRNA or sgRNA based editing, after 2-3 days of recovery from transfection, cells are plated on 10cm dishes at the same density for colony formation without any drug selection.

**Production of rescue cells**

cDNA sequences for bovine VPS50-54 subunits were amplified from total cDNA reverse transcribed from wt MDBK cells and cloned into a PiggyBac vector after a CAG promoter followed by a Puromycin selection cassette connected via an IRES sequence. Representative VPS50-54 KO clones were then transfected with the cDNA expression vectors and pCMV-hypBase using Neon . Cells were then selected with Neomycin 48 hours after transfection and expanded for experiments.

**BHV-1 infection**

For the screening, 10^6^ cells were plated in each T175 flask (9 flasks per repeat) 12 hours prior to infection (passage 4 or 5 with 9-10 days of puro selection and culture post transduction). Cells in each flask were infected with GFP tagged BHV-1 at a MOI=2 in media containing 2% horse serum. After one hour	the virus inoculum was removed and replaced	with 20ml of fresh media with 2% Horse Serum. This time point was counted as 0-hour post infection (0hpi). BHV-1 infections for single gene studies are scaled down with the quantities of viruses calculated based on the desired MOI. Where Phosphonoacetic acid (PAA) was used, it was added to the media 10 minutes before infection with at the final concentration of 200 µg/mL.

**Plaque assays**

One day before plaquing, approximately 5 x10^5^ cells from each cell line are plated in 2ml culture media with 2% horse serum per well on 6-well plates. One day later, cells were infected with 10-fold serial dilutions of wt or GFP tagged BHV-1 virus in 1ml of media with 2% horse serum. After one hour the virus inoculum was removed and r 2mls of Avicel overlay (culture media with 2% horse serum and 0.5% Avicel) was added to each well. The plates are returned to the incubator, and four days later, cells were fixed by adding 1ml 10% Neutral Buffered Formalin. After one hour of incubation at room temperature, the liquid mix is removed and 1ml 0.1% Toluidine Blue is added to each well. After staining for one hour at room temperature, the wells are washed gently with tap water and inverted to dry overnight. Plaque numbers are counted under an inverted microscope and the plates are also scanned using an Epson document scanner. The scanned images were processed by ImageJ to count and measure sizes of plaques.

**Statistical analysis**

Unless otherwise stated, all statistical pairwise comparisons between two series of samples were done using the “Analysis Tool ANOVA: Single Factor” provided by Microsoft Excel with Version 16.34 (20020900), or student t-test for pairwise comparison of two means.

**FACS sort**

To prepare cells for sort, media was removed carefully at the appropriate time point without dislodging cells and 1.5ml 0.5% Trypsin was added to the flasks immediately without PBS wash. After 5 minutes of incubation at 37° Celsius, trypsin was neutralized with 8ml of culture media and the cell suspension harvested. The flasks were t rinsed with 10mls of PBS and trypsinized again for 3 minutes to recover the residual cells. All cell suspensions from the two rounds of trypsinization and the PBS washes were combined, centrifuged at 1200g for 5 minutes, washed once with 50ml of PBS and recentrifuged. The cell pellet was then resuspended in 8mls of PBS containing 2mM EDTA. The cell suspension was kept on ice and filtered with a 0.22um nylon mesh prior to sort and maintained at 6°C during sort. Four fractions of live cells, GFP negative, ~10% GFP low, ~10% GFP Medium, and ~10% GFP High cells are collected. Collection tubes are pre-filled with 200ul horse serum prior to sort.

**Genomic DNA isolation and PCR for NextSeq**

After FACS sort, collected cells were centrifuged at 1200g for 5 minutes, washed once with PBS and the pellet stored at -80° Celsius until genomic DNA isolation. Based on the number of cells recovered, genomic DNA was isolated from the samples using one of the following kits: Quick-DNA Microprep kit, Quick-DNA Miniprep Plus Kit, DNeasy Blood & Tissue kit, NucleoSpin Blood L Column, or NucleoSpin Blood XL Column. For non-infected control samples, fragment enrichment by HindIII-HF digest and gel extraction was conducted to reduce the number of PCR retractions required to prepare samples for Illumina sequencing. Briefly, 300ug of genomic DNA was digested with 2,000 units of HindIII-HF overnight for 20 hours and resolved on a 0.7% Agarose gel. The fragment containing the hU6-sgRNA cassette is 1485bps, a gel slice containing DNA in the range of 1.2kb -1.8kb was excised from the gel and the DNA extracted from the slice using the NucleoSpin Gel and PCR Clean-up Kit.

To prepare samples for NextSeq, 1- 2ug genomic DNA or 0.5-1ug enriched genomic DNA fragments were used as template in the 1^st^ PCR per reaction, with limited cycles for faithful representation of the original gRNA distributions in the samples. The PCR was performed using Q5 Hot Start polymerase and a cocktail of ten forward primers containing the staggering sequences (primers PCR_F1 to PCR1_F10, Supplementary Table S7) and the reverse primer PCR1_R (Supplementary Table S7). For every sample, 5ul from each PCR reaction was pooled and cleaned up using a PCR cleanup column. The cleaned-up product was then used in a 2^nd^ PCR with 10 cycles for barcoding, using NEBNext Ultra II Q5 polymerase and a unique combination of one forward primer idx_Sxxx (Supplementary Table S7) and one reverse primer idx_Nxxx (Supplementary Table S7). The 2^nd^ PCR reaction was then cleaned up using 1 Volume AMPure XP beads on a magnetic stand and eluted with EB buffer from a Qiagen miniprep kit. The purified 2nd PCR product is quantified using a Qubit dsDNA HS Assay Kit and a Qubit 2.0 machine. The products were also visualized, and quality confirmed by running 2ul of each purified product on a 2% Agarose gel. Purified products for all samples were pooled into	and sent	for sequencing on a NextSeq 500 machine. The sample was sequenced using a Single Read 75bp High Output Kit v2 with 25% PhiX spike in, resulting in ~320 million reads passing quality control. The reads are then demultiplexed and assigned to each sample based on their unique barcode combinations.

All illumine sequencing was carried out on a Nextseq 500 machine at the Edinburgh Clinical Research Facility (ECRF) using NextSeq 500/550 High Output Kit v2.5 (75 Cycles). A spike-in of 25% PhiX was added to the sample mix to increase sequence diversity.

**NextSeq reads processing and data analysis**

For each sample, the sequencing reads were checked for quality using FastQC v0.11.4 and trimmed using cutadapt v1.16 from the 5’ until sequence CGAAACACCG (inclusive) and from the 3’ until GTTTAAGAGC (inclusive), discarding all untrimmed sequences and keeping only trimmed reads with 20bp in length. The copy number of trimmed reads matching each guide in the CRISPR library was then counted using the *count* sub-command from the MAGeCK package v 0.5.8^9^. Pairwise comparisons of guide copy numbers between samples were done using the *test* sub-command from MAGeCK. Candidate genes with significantly depleted or enriched guides were selected based on a cut-off of adjusted P-Value < 0.005, FDR<0.1 and l2fc>=1 or <=-1.

**Gene Ontology analysis**

All GO enrichment analyses were completed using PANTHER Overrepresentation Test (Released 20190711) and GO Ontology database (Released on 2019-07-03) hosted on website <http://geneontology.org/>. The queries were conducted using FISHER as the Test Type and all genes in database for Homo sapiens as Reference List. We chose the Homo sapiens list instead of that for Bos Taurus due to better data availability.

**Viral genomic DNA sample collection and quantification**

Following infection, the supernatant and cell pellet samples were harvested at time points 0, 2, 4, 6, 8, 24, 48 and 72 h post infection. For nucleus and cytoplasm separation, cell pellets were gently resuspended with 1 × Hypotonic buffer (20 mM Tris-HCl, Ph 7.4; 10 mM NaCl; 3 mM MgCl2) on ice for 15 minutes, and then added detergent (10% NP40) and vortexed. After centrifuge for 10 min at 3,000 rpm at 4 °C, the supernatant contained the cytoplasmic fraction were transferred and saved. Then the nuclear fraction pellet was resuspended with RLT buffer according to the protocol of RNeasy Plus Mini Kit (Qiagen, Hilden, Germany). Viral DNA copies were determined by qPCR, and the standard curves of UL23 gene and host gene PVRL2 were used. As a negative control, a final concentration of 200 µg/mL PAA was added to the media 10 minutes before infection with BHV-1. qPCR was conducted using [LightCycler® 480 SYBR Green I Master](https://shop.roche.com/shop/store/ProductDisplay?catalogId=10001&partNumber=3.5.8.1.1.2) reagents and samples were run in a LightCycler 480 machine.

**Reverse transcription and Quantitative PCR**

To compare the viral mRNA levels by reverse transcription and qPCR, total RNA from cells was isolated using the RNeasy Plus Mini Kit (Qiagen, Hilden, Germany). Then RNA was DNase-treated, and cDNA was synthesized using a QuantiTect reverse transcription kit (Qiagen, Hilden, Germany). Quantitative real-time RT-PCR was carried out using the LightCycler 480 System and LC480 SYBR Green 1 Master (Roche) following the protocol provided by the manufacturer. Relative bovine ICP4 (bICP4), UL23 and UL35 mRNA levels were quantified by qPCR using 18S rRNA internal controls to normalize template input and presented as fold change against the CTRL cells. Gene differential expression between samples was calculated using 2-ΔΔCT method.

**VP26-GFP growth tracing**

30,000 cells were seeded in RPMI media without Pheno red plus 5% HS on a 96-well clear bottom dark walled culture plate 12 hours prior to reading. For each cell line, four replicates were set up. The outside wells were avoided to reduce bias from evaporation. Cells were infected by the GFP tagged BHV-1 virus either at MOI=3 or MOI=0.1 and the first reading was taken at 0 h.p.i. The cells were incubated in 5% CO2 at 37 °C while the Fluorescent Intensity (FI) from each well was recorded by CLARIOstar Plus Microplate Reader (BMG LabTech) every 10 minutes for 72 hours.

**STRING clustering analysis**

STRING analysis of lists of candidate genes were queried using the “Multiple Proteins by Names/Identifiers” search box against the database for Bos Taurus on: <https://string-db.org/cgi/network.pl?taskId=4414a1leHqHx>.

Default values were used for all settings except the following: meaning of network edges = molecular action; Clustering method = MLC clustering.

**Table S3. Plasmids used in the study**

| **Plasmid name** | **Source** | **Purpose** |
| --- | --- | --- |
| pKLV-U6gRNA(BbsI)-PGKpuro2ABFP | Gift from Kosuke Yusa | Lentivirus g2 Library cloning |
| pCMV-hypBase | Gift from Kosuke Yusa | Piggyback transposase |
| pKLV2-U6gRNA5(BbsI)-PGKpuro2ABFP-W | Addgene # 67974 | Lentivirus g5 library cloning |
| lentiGuide-Puro | Addgene # 52963 | Lenti delivery of sgRNA |
| pSpCas9(BB)-2A-GFP (PX458) | Addgene # 48138 | CRISPR/Cas9 expression |
| PMD2.G | Addgene # 12259 | lentivirus packaging |
| psPAX2 | Addgene # 12260 | lentivirus packaging |
| pHIV-EGFP | Addgene # 21373 | Control lentivirus |
| pMT2.0_bovRosa26_EF1a_Cas9_blast | this study | Cas9 targeting to rosa26 |
| PB_U6gRNA2-PGKpuro2ABFP-W | this study | PiggyBac g2 Library cloning |
| PB_U6gRNA5-PGKpuro2ABFP-W | this study | PiggyBac g5 library cloning |
| PB_U6gRNA2-CAGpuro | this study | PiggyBac delivery of sgRNA |
| PMC-DeltaTAL | this study | TALEN assembly |
| PMC-bovRosa26_1.6L | this study | Rosa26 targeting |
| PMC-bovRosa26_1.6R | this study | Rosa26 targeting |
| PMC-bovTRIM5a_2.2L | this study | TRIM5a targeting |
| PMC-bovTRIM5a_2.2R | this study | TRIM5a targeting |
| PB_VPS51R | this study | cDNA overexpression |
| PB_VPS52R | this study | cDNA overexpression |
| PB_VPS53R | this study | cDNA overexpression |

**Note:** Plasmids generated in this study are available upon request.

**Table S4. Individual CRISPRS and TALENs used in this study**

| **Nuclease name** | **Sequence** | **Purpose** |
| --- | --- | --- |
| Rosa26_1.6L | NN NN NN HD HD NG NI NN NI NI NN NI NI NG HD HD HD NG | Rosa26 targeting |
| Rosa26_1.6R | HD NN NI HD NI NG NN NN NI NN NN HD NN NI NG NN NI HD NN NI | Rosa26 targeting |
| TRIM5_1L | NI NN NN NI NN NG NN HD NI NI NI NG NN NG NG NN | TRIM5 KO |
| TRIM5_1R | NG HD NG HD NI HD NG NG NN NG HD NG HD NG NN | TRIM5 KO |
| VPS50_sgRNA | AGGGACCCGGAGACTCTCAA | KO |
| VPS51_sgRNA | GTTGTAGTTCTCATAGACCA | KO |
| VPS52_sgRNA | GGAGCTTGTTGATGGTCTCG | KO |
| VPS53_sgRNA | GTTCAGCGTTGTGATCGAGG | KO |
| VPS54_sgRNA | AGGCTGAGTCTGGTGTACAG | KO |
| CTNNB1 g1 | GAGCGGTAAAGGCAATCCTG | KO |
| CTNNB1 g2 | GTTCCCTGAGACGCTAGATG | KO |
| MAPK8 g1 | GTAGTAGCGAGTCACTACGTA | KO |
| MAPK8 g2 | GTCGCTACTACAGAGCACCTG | KO |
| MAPK9 g3 | GACCCTGAAGATCCTCGACTT | KO |
| MAPK9 g4 | GTGGTGCACGCTGTACGAGCC | KO |
| Oct-1 g1 | GCTGGCGGAATGCTGCTGCA | KO |
| Oct-1 g2 | GCTGTATGGGCTGAGACAAG | KO |
| SGK1 g1 | GTGCAGGTAACCCAAGGCAC | KO |
| SGK2 g1 | GCAGCATAGAATCGAGCCCG | KO |
| SGK3 g1 | GTTGTTCTACCATCTCCAGA | KO |

**Table S5. Primers used for cloning, gene targeting and qPCR**

| **Primer name** | **Sequence** | **Purpose** |
| --- | --- | --- |
| 79-mer-U1 | GCAGATGGCTCTTTGTCCTA | g2, g5 library cloning |
| 79-mer-L1 | GCGACGAGAAGACTGTAAAAC | g2 library cloning |
| 79-mer-L1v2 | GCGACGAGAAGACTAAAAC | g5 library cloning |
| btRosa26_NJ_F1 | CGCTGCCTGAAGGACAAGAC | T7/TIDE |
| btRosa26_NJ_R1 | CGGTGTAGCAACGGTCTCAAA | T7/TIDE |
| btRosa26_4k_F1 | TCGTGAGGGTAGGTCTCTCTT | Rosa26 targeting |
| btRosa26_4k_R1 | ACCTGCCAACCAACCAACC | Rosa26 targeting |
| TRIM5a NJ F7 | CCACCCTATTCTCATCATCT | T7/TIDE |
| TRIM5a NJ R7 | GTTGATATACTGAAAAAACATTTG | T7/TIDE |
| VPS50_NJ_F | ATGGTCTCGGTGAAAGAAGGGGAA | T7/TIDE |
| VPS50_NJ_R | CAAGCCCCAAGCCTATTGCATTATA | T7/TIDE |
| VPS51_NJ_F | CCTGGTAACTCTTGAGCTATGCCAC | T7/TIDE |
| VPS51_NJ_R | ATTTCTTTTTAAAATTTCAATGTCT | T7/TIDE |
| VPS52_NJ_F | GTTTTTGTTGTTGCTGCCGTGACTA | T7/TIDE |
| VPS52_NJ_R | TCACGCCTGCCCATCTACTTTCTAT | T7/TIDE |
| VPS53_NJ_F | CGGGTAAAGTCTGCTGAAGAAATGC | T7/TIDE |
| VPS53_NJ_R | TGCCGTTTCCTGCGAGTCATTCAT | T7/TIDE |
| VPS54_NJ_F | TGCTCACCAGATCTCTTTACGTTCA | T7/TIDE |
| VPS54_NJ_R | ACGACTGAGCAACTAAACGGAACTG | T7/TIDE |
| CTNNB1_NJ_F2 | TCCTGGTAGTAATATTGATGCTGT | T7/TIDE |
| CTNNB1_NJ_R2 | TGTCTCTACTTACCTGGTCCTC | T7/TIDE |
| MAPK8_NJ_F2 | TGCCATCAAGCTAATTTCTCAGAT | T7/TIDE |
| MAPK8_NJ_R2 | GCGGGCCCACTATACACTTC | T7/TIDE |
| MAPK9_NJ_F2 | ACCGGAATGTTGGGAACTGA | T7/TIDE |
| MAPK9_NJ_R2 | CTCATCTGTGAACCATGATAATCTA | T7/TIDE |
| Oct-1_NJ_F1 | TGCCATATGAAGTTGGGTAGC | T7/TIDE |
| Oct-1_NJ_R1 | CCTCTGCTCCAGAAATACGC | T7/TIDE |
| SGK1_NJ_F2 | CAGACGGCCGACAAACTGTA | T7/TIDE |
| SGK1_NJ_R2 | AAGCTTGGCCTATGGTCTCC | T7/TIDE |
| SGK2_NJ_F1 | CTGCCTGTAAGTCCTCCGAC | T7/TIDE |
| SGK2_NJ_R1 | TGGGGATCTCACGATGGAGT | T7/TIDE |
| SGK3_NJ_F2 | CCGCCTCTTCTAAGGAGCTTT | T7/TIDE |
| SGK3_NJ_R2 | GAGTACTTCCCAAGTGCCTCC | T7/TIDE |
| 18S_qPCR_F | TGTGATGCCCTTAGATGTCC | Host cell gDNA qPCR |
| 18S_qPCR_R | TTATGACCCGCACTTACTGG | Host cell gDNA qPCR |
| bICP4_qPCR_F | CGGAGAGCAGCGAGGACGACGG | ICP4 cDNA qPCR |
| bICP4_qPCR_R | GCTTCGATGGCGGCGGCTATGA | ICP4 cDNA qPCR |
| UL35_qPCR_F | GCAGATTTTGCATGTGCTAAACGCC | UL35 cDNA qPCR |
| UL35_qPCR_R | CGTGGTCGTAGGTCGCAAACAT | UL35 cDNA qPCR |
| UL23_qPCR_F | CTCTGCTACCCCTTCGCCCGCTACT | UL23 cDNA qPCR |
| UL23_qPCR_R | AGGGTGCACACGACGAGGTTGGC | UL23 cDNA qPCR |
| PVRL2_qPCR_F | AGCCAGAAAGAATCTAAGGGCCAGG | PVRL2 gDNA qPCR |
| PVRL2_qPCR_R | CCCAGGTCGTCCGAATTATCATTCT | PVRL2 gDNA qPCR |

**Note:** iVT = *in vitro* transcription;

**Table S7. Primers used for NextSeq**

| **Primer** | **Sequence** |
| --- | --- |
| PCR1_F1 | TCGTCGGCAGCGTCAGATGTGTATAAGAGACAGTCTTGTGGAAAGGACGAAACACCG |
| PCR1_F2 | TCGTCGGCAGCGTCAGATGTGTATAAGAGACAGATCTTGTGGAAAGGACGAAACACCG |
| PCR1_F3 | TCGTCGGCAGCGTCAGATGTGTATAAGAGACAGGATCTTGTGGAAAGGACGAAACACCG |
| PCR1_F4 | TCGTCGGCAGCGTCAGATGTGTATAAGAGACAGCGATCTTGTGGAAAGGACGAAACACCG |
| PCR1_F5 | TCGTCGGCAGCGTCAGATGTGTATAAGAGACAGTCGATCTTGTGGAAAGGACGAAACACCG |
| PCR1_F6 | TCGTCGGCAGCGTCAGATGTGTATAAGAGACAGATCGATCTTGTGGAAAGGACGAAACACCG |
| PCR1_F7 | TCGTCGGCAGCGTCAGATGTGTATAAGAGACAGGATCGATCTTGTGGAAAGGACGAAACACCG |
| PCR1_F8 | TCGTCGGCAGCGTCAGATGTGTATAAGAGACAGCGATCGATCTTGTGGAAAGGACGAAACACCG |
| PCR1_F9 | TCGTCGGCAGCGTCAGATGTGTATAAGAGACAGACGATCGATCTTGTGGAAAGGACGAAACACCG |
| PCR1_F10 | TCGTCGGCAGCGTCAGATGTGTATAAGAGACAGTACGATCGATCTTGTGGAAAGGACGAAACACCG |
| PCR1_R | GTCTCGTGGGCTCGGAGATGTGTATAAGAGACAGCTAAAGCGCATGCTCCAGAC |
| idx_S502 | AATGATACGGCGACCACCGAGATCTACACCTCTCTATTCGTCGGCAGCGT*C |
| idx_S503 | AATGATACGGCGACCACCGAGATCTACACTATCCTCTTCGTCGGCAGCGT*C |
| idx_S504 | AATGATACGGCGACCACCGAGATCTACACAGAGTAGATCGTCGGCAGCGT*C |
| idx_S517 | AATGATACGGCGACCACCGAGATCTACACGCGTAAGATCGTCGGCAGCGT*C |
| idx_N701 | CAAGCAGAAGACGGCATACGAGATTCGCCTTAGTCTCGTGGGCTCG*G |
| idx_N702 | CAAGCAGAAGACGGCATACGAGATCTAGTACGGTCTCGTGGGCTCG*G |
| idx_N703 | CAAGCAGAAGACGGCATACGAGATTTCTGCCTGTCTCGTGGGCTCG*G |
| idx_N704 | CAAGCAGAAGACGGCATACGAGATGCTCAGGAGTCTCGTGGGCTCG*G |
| idx_N705 | CAAGCAGAAGACGGCATACGAGATAGGAGTCCGTCTCGTGGGCTCG*G |
| idx_N706 | CAAGCAGAAGACGGCATACGAGATCATGCCTAGTCTCGTGGGCTCG*G |

**Note:** 1^st^ PCR primers (PCR1_F1 to PCR1_R) were ordered as regular oligos and index primers (idx_S502 to idx_N706) as Ultramers from IDT with the last base (*) Phosphorothioated to prevent degradation. The barcodes are underlined.

**Table S8. Reagents and consumables used in the study**

| **Reagent** | **Supplier** | **Cat. I.D.** | **Purpose** |
| --- | --- | --- | --- |
| 2XHBS | VWR | J62623.AK | Lentivirus packaging |
| 2.5M CaCl_2_ | Jena Bioscience | BU-103-JEN | Lentivirus packaging |
| DNase/RNase-Free Distilled Water | Life Technologies | 10977035 | Various |
| DMEM | Sigma | D5796 | Tissue culture |
| Horse serum | Sigma | H1138 | MBDK culture |
| Horse serum | Gibco | 26050088 | MBDK culture |
| Bovine Serum | Gibco | 10270 | HEK293FT culture |
| Fetal bovine serum | Gibco | 16170-078 | HEK293FT culture |
| Pen/Strep | Gibco | 15140122 | Cell culture |
| NEAA | Gibco | 11140035 | MBDK culture |
| Sodium Pyruvate | Gibco | 11360039 | MBDK culture |
| L-Glutamine | Gibco | 25030024 | MBDK culture |
| Trypsin solution | Sigma | T3924-100ML | Cell culture |
| BloodStor 100 DMSO | StemCell Technologies | 07951 | Cell freezing media |
| Blasticidin S HCl | Corning B.V. | 30-100-RB | Rosa26 targeting |
| Blasticidine S hydrochloride | Sigma | 15205-25mg | Rosa26 targeting |
| G418 | Sigma | G8168-10ML | HEK293FT culture |
| Puromycin Dihydrochloride | ThermoFisher | A1113803 | Library transduction |
| Saline, Buffered; HEPES; 1M | GE Hyclone | 10204932 | Library packaging |
| 150mm Petri dish | Corning | 10117320 | Library packaging |
| Syringe Filter, PES, 0.45um | Fisherbrand | 15216869 | Lentivirus packaging |
| 0.45um filter, 500ml | Corning | 430770 | Lentivirus filtration |
| Polybrene | Merck | TR-1003-G | Lentivirus transduction |
| NEB PCR Cloning Kit | NEB | E1203S | Construct cloning |
| Zero Blunt TOPO Cloning Kit | ThermoFisher | 450245 | PCR cloning |
| 10-betaStable Outgrowth Medium | NEB | B9035S | Library cloning |
| Ampicillin sodium salt | Sigma | A9518-25G | Library cloning |
| BbsI-HF | NEB | R3539S | Library cloning |
| BpiI(BbsI) (10U/uL) | Thermo Scientific | ER1011 | Library cloning |
| Quick Ligation Kit | NEB | M2200L | Library cloning |
| Adenosine 5’-Triphosphate (ATP) | NEB | P0756S | Library cloning |
| QIAquick Nucleotide Removal Kit | Qiagen | 28304 | Library cloning |
| HindIII-HF | NEB | R3104M | Genomic DNA digestion |
| NEBNext Ultra II Q5 Master Mix | NEB | M0544L | 2^nd^ PCR for NextSeq |
| 2X Q5 Hot Start Master Mix | NEB | M0494L | 1^st^ PCR for NextSeq |
| 2X Phusion HF Master Mix | NEB | M0531L | PCR for T7 and Sanger seq |
| 2X DreamTaq Green PCR Master Mix | Thermo Scientific | K1081 | Routine PCR |
| PrimeSTAR GXL DNA polymerase | Takara Bio | R050A | Long Range PCR |
| NEBuilder HiFi DNA Assembly Cloning Kit | NEB | E5520S | Construct cloning |
| Plasmid plus Midi kit | Qiagen | 12943 | Plasmid prep |
| Plasmid plus Maxi kit | Qiagen | 12963 | Library plasmid prep |
| Qiaprep Spin Miniprep kit | Qiagen | 27104 | Plasmid preparation |
| NucleoSpin Blood L Column | Machery-Nagel | 12761021 | Genomic DNA isolation |
| NucleoSpin Blood XL Column | Machery-Nagel | 12731021 | Genomic DNA isolation |
| Ribonuclease A | Sigma | R6513-1G | Genomic DNA isolation |
| DNeasy Blood & Tissue kit | Qiagen | 69504 | Genomic DNA isolation |
| Quick-DNA Microprep kit | Zymo Research | D3020 | Genomic DNA isolation |
| Quick-DNA Miniprep Plus Kit | Zymo Research | D4068 | Genomic DNA isolation |
| Qubit dsDNA HS Assay Kit | Invitrogen | Q32854 | DNA quantification |
| QuickExtract DNA Extraction Solution | Lucigen | QE09050 | Genomic DNA extraction |
| 10-beta Electrocompetent E Coli | NEB | C3020K | Library cloning |
| Gene Pulser/MicroPulser Cuvettes, 0.1cm gap | Bio-rad | 1652089 | Library cloning |
| NEB Stable Competent Cells | NEB | C3040I | Construct cloning |
| Top10 Competent Cells | Invitrogen | C404003 | Plasmid transformation |
| Agencourt AMPure XP | Beckman Coulter | A63880 | PCR cleanup for NextSeq |
| Agarose UltraPure | Life Technologies | 16500500 | DNA/RNA electrophoresis |
| ChargeSwitch PCR Clean-up kit | Invitrogen | CS12000 | PCR cleanup for Sanger Seq |
| Zymoclean Gel DNA Recovery | Zymo Research | D4001 | Library cloning |
| NucleoSpin Gel and PCR Clean-up Kit | Machery-Nagel | 11992242 | Gel extraction and PCR cleanup |
| MinElute PCR Purification Kit | Qiagen | 28004 | PCR cleanup |
| Neon Transfection System 100 uL Kit | ThermoFisher | MPK10025 | MDBK transfection |
| T7 Endonuclease I | NEB | M0302S | Gene editing detection |
| HiScribe T7 Quick High Yield RNA synthesis kit | NEB | E2050s | sgRNA *in vitro* transcription |
| Hiscribe T7 ARCA mRNA Kit | NEB | E2060S | mRNA *in vitro* transcription |
| RNASecure RNase Inactivation Reagent | ThermoFisher | AM7005 | RNA treatment |
| RNeasy mini Kit | Qiagen | 74104 | RNA isolation and cleanup |
| Doxycycline hyclate | Sigma | D9891-1G | dCas9 induction |
| Avicel RC/CL™ | FMC Biopolymer | RC-581 | Plaque assays |
| 10% Neutral Buffered Formalin | CellPath | BAF-0010-20A | Plaque assays |
| Toluidine Blue | Sigma | T3260 - 100G | Plaque assays |
| [LightCycler® 480 SYBR Green I Master](https://shop.roche.com/shop/store/ProductDisplay?catalogId=10001&partNumber=3.5.8.1.1.2) | Roche | 04887352001 | qPCR |

2.	*Protocol for Lentivial Vector (LV) Production (2 nd Generation Packaging)*.

3.	Cermak, T. *et al.* Efficient design and assembly of custom TALEN and other TAL effector-based constructs for DNA targeting. doi:10.1093/nar/gkr218
